# Engineering nanocondensate formation through sequence composition and patterning

**DOI:** 10.64898/2026.02.17.706365

**Authors:** Timo N. Schneider, Florence Stoffel, Marco A. Bühler, Karolína Mrzílková, Milad Radiom, Paolo Arosio

## Abstract

Many biological proteins can assemble into dynamic, non-stoichiometric structures known as biomolecular condensates. Although typically observed at micrometer scales in vitro, recent evidence shows that these condensates often appear as nanoscale assemblies both in vitro and in cells. Moreover, biochemical reactions can be more efficiently promoted in nanoscale condensates than in micron-sized droplets, due to mass-transfer limitations and interfacial effects. Therefore, in analogy with colloids, the function of condensate materials can be engineered by tuning their size distribution. However, controlling the size of condensates remains challenging, as the molecular mechanisms that prevent small condensates from coarsening into larger ones are still poorly understood. Here, we developed and applied a computational pipeline that combines high-throughput molecular simulations, machine learning, and mixed-integer linear programming to design phase-separating peptides that form metastable nanocondensates across a broad range of experimental conditions. In addition to experimentally validating these peptides, we elucidate the underlying molecular mechanisms and derive initial design rules. In particular, we show that scaffold net charge combined with sequence blockiness can lead to high phase separation propensity while simultaneously yielding low interfacial tension, thereby slowing ripening. Moreover, these combined properties induce an electrostatics-driven alignment of molecules at the interface, which generates an additional size-dependent coalescence barrier. We further show that these features are shared by biological proteins, providing a possible mechanistic basis for the widespread occurrence of nanocondensates in biological systems. Altogether, our findings shed light on the molecular mechanisms behind nanocondensate formation, and provide a platform to design nanocondensates for several potential applications in bioengineering and biotechnology.

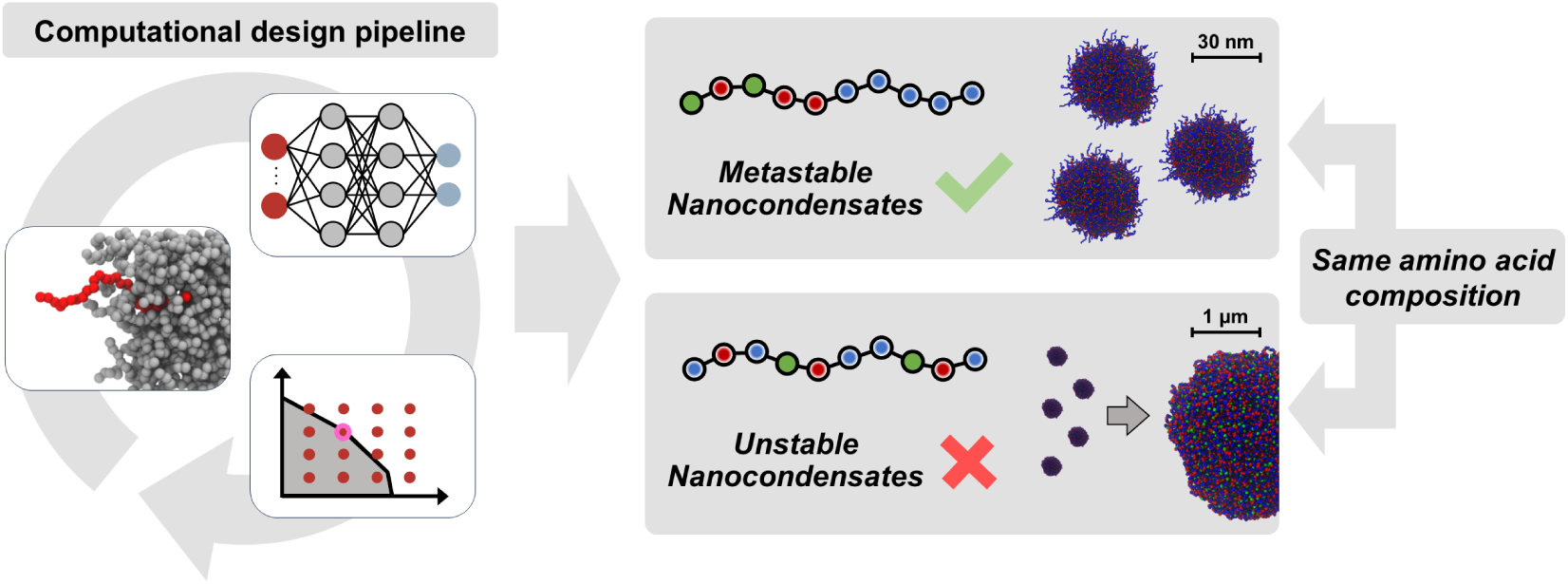

## Introduction

In addition to membrane-bound compartments, cells can regulate their metabolism in space and time through the formation of membraneless protein- and RNA-rich compartments, known as biomolecular condensates.^1–4^ In the last decade, many studies investigated these condensates at the micron scale, largely based on microscopy techniques. However, an increasing number of recent findings have revealed the presence of mesoscale assemblies in the ∼100 nm range for a variety of phase-separating proteins,^5–7^ including *α*-synuclein,^8^ FET family proteins,^9–11^ and tau,^12^ which have been indicated as nanocondensates, mesoscopic pre-percolation clusters, or nanoclusters. These assemblies share many properties with micron-scale condensates, such as molecular exchange with the surrounding, lack of fixed stoichiometry, stimuli-responsiveness, and reversibility of their formation, and have been observed both below and above the critical concentration for macroscopic phase separation.^9,10,13–16^ In contrast to other types of nanoscale molecular assemblies, such as stoichiometric protein oligomers, micelles, or microphases,^17^ nanocondensates are observed in broad, often heavy-tailed size distributions, and their dimensions can exceed those of the scaffold molecule by orders of magnitude.^16^ A recent study has shown that a substantial fraction of the proteome is organized within this size range of ∼100 nm, making nanocondensates a highly relevant target of investigation.^18^

Despite their presence in many *in vitro* and *in vivo* systems, the mechanisms underlying nanocondensate formation and their colloidal stability remain unclear. In a simple phase-separating system, such structures are expected to be unstable, as small assemblies tend to grow into a single large dense phase via Ostwald ripening and coalescence events, to minimize total interfacial area and, consequently, the free energy of the system. Moreover, nanocondensates do not represent prenucleation clusters predicted by classical nucleation theory, since these structures should be much smaller and consist of only a few molecules.^9,10^ Nanocondensates could represent thermodynamically stable species at equilibrium. For example, theoretical and simulation studies show that a charge imbalance within the nanocon-densate giving rise to long-range repulsion combined with short-range attraction can lead to nanocondensates being more stable than a macroscopic dense phase. ^19,20^ In addition, lattice-based simulations have suggested that copolymers composed of monomers with different interaction strengths can give rise to nanocondensate formation.^10^ According to another hypothesis, molecules can adopt distinct conformations at the nanocondensate interface, exposing more hydrophilic regions to the dilute phase. This results in ultra-low interfacial tension, giving rise to an equilibrium distribution that extends into the mesoscopic size range.^14,21^

Another possibility is that nanocondensates represent kinetically trapped, metastable species for which ripening and coalescence events are inhibited. Indeed, in analogy to emulsions stabilized by surfactants, the size distribution of condensates can be modulated by accumulation of molecules or particles at the interface.^22–25^ Kinetic stabilization of nanocondensates was observed also in single-protein systems.^8–10^ In these cases, the scaffold net charge and the zeta potential of condensates play a key role in modulating coalescence, in analogy with colloidal systems.^26–29^ Furthermore, saturation of valences of scaffold molecules can also inhibit coalescence.^30,31^ Finally, the coupling of phase separation with active processes has also been shown to provide an additional mechanism to maintain condensates out of equilibrium and inhibit Ostwald ripening.^32,33^

Irrespective of their mechanism of formation, these dynamic nano-materials are highly attractive for nanotechnology and material science. For instance, nanocondensates can accelerate biochemical reactions even more effectively than micron-scale condensates due to reduced mass transport limitations.^34–36^ Moreover, the interface of the condensates has peculiar properties and can mediate biochemical reactions and aggregation events.^8,37–40^ Since these interfacial effects are more prominent at the nanoscale, in analogy with colloids, it is thus possible to design condensates with different emergent properties simply by controlling their size distribution.

In this study, we de novo design phase-separating peptides forming kinetically arrested nanocondensates that resist ripening and coalescence. Because the intermolecular interactions that drive phase separation also lead to an energetic cost for interface formation, systems with a strong tendency to demix typically exhibit high interfacial tension *γ*.^41–43^ Consistent with this correlation, the introduction of molecules that increases the propensity for phase separation also often increases interfacial tension.^44,45^ Here, we aimed to design peptides that break this correlation, yielding single-component peptide coacervates with minimal interfacial tension and thus slow Ostwald ripening, while simultaneously maintaining a strong driving force for phase separation to incorporate a large fraction of the monomers into nanocondensates (Figure 1A).

**Figure 1:**
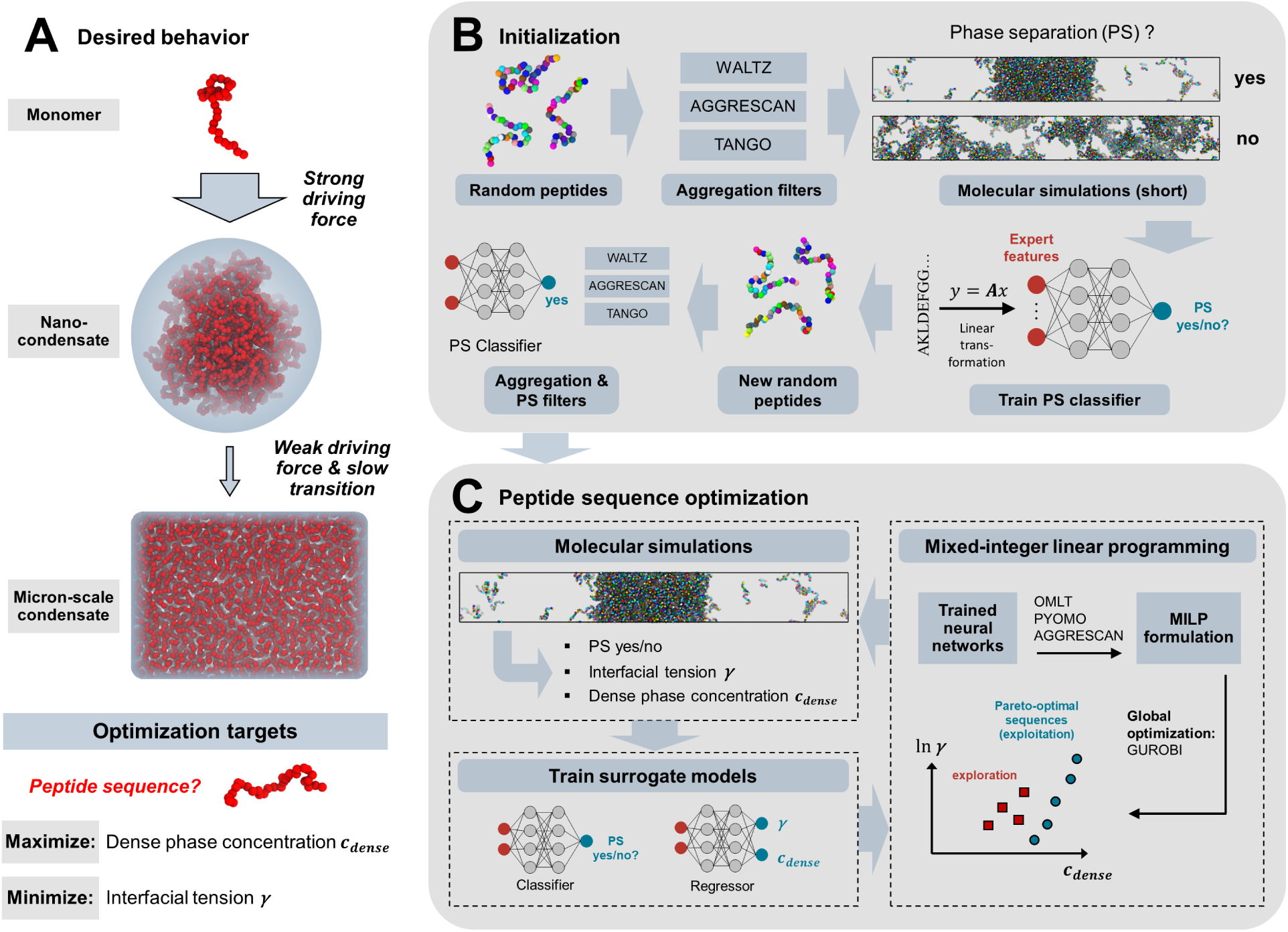
Design strategy to create peptides forming nanocondensates. (A) The core idea at the basis of this work is the identification of peptides with a high propensity to phase separate (high *c*_dense_) into condensates characterized by low interfacial tension *γ*. This combination provides a strong driving force for nanocondensate formation but only a weak driving force for growth into micron-scale condensates, and inhibits Ostwald ripening. (B) Fully random peptides were tested for phase separation using coarse-grained simulations, and an initial classifier was first trained and then used to exclude non–phase separating peptides. (C) The optimization loop started with molecular simulations that, in the case of phase separation, quantified *c*_dense_ and *γ*, after which the classifier and regressor were updated with the new simulation data. The optimization problem was formulated as an MILP, which was used to identify new sequences to simulate in the next iteration.

We designed and experimentally validated such peptides, combining high-throughput coarse-grained simulations with an active learning algorithm, the latter building on a recently developed approach.^25^ Our results show that the decoupling between phase separation and interfacial tension is effectively mediated by sequence patterning in addition to net sequence charge. At the molecular level, we demonstrate that the metastability of these nanocondensates originate from a size-dependent interfacial structuring of the peptide coacervates induced by the interphase electric potential,^38,46^ which reduces interfacial tension and also inhibits coalescence. We further show that insights from these minimal peptides are shared by natural protein systems and that the mechanisms highlighted by this study combine previously proposed explanations for the formation of nanocondensates.

## Results and Discussion

### Identification of phase separation-prone peptides leading to low interfacial tension

We aimed at designing peptides with low interfacial tension (*γ*) and low dilute phase concentration (*c_∞_*) which inhibits the rate of Ostwald ripening (*ω*) according to Lifshitz, Slezov, and Wagner theory:^47,48^

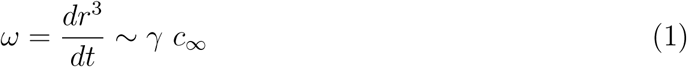

where *r* refers to the number average condensate radius. We did not explicitly include coalescence in the design loop because it is more challenging to quantify in a high-throughput in silico setting. We instead investigated coalescence behavior in a second step for a smaller set of selected candidates. Moreover, instead of directly minimizing *c_∞_*, we maximized the dense phase concentration *c*_dense_, which is anticorrelated with *c_∞_* and computationally less expensive to measure.

We fixed the peptide length at 30 AA to ensure accessibility for chemical synthesis, while still leading to a vast design space of 20^30^ possibilities, requiring efficient quantification of objectives as well as navigation of this space. High-throughput molecular simulations, potentially combined with active learning, have been proven efficient for designing polymers, peptides, and disordered proteins for a variety of design targets.^25,49–58^ We kept the optimization unbiased by starting with fully random peptides, where each of the 20 amino acids was uniformly sampled (Figure 1B). To avoid aggregation-prone regions, we filtered the peptides using the aggregation predictors Waltz,^59^ TANGO,^60^ and AGGRESCAN.^61^ This led to 500 sequences for which short coexistence simulations were run using the one-bead-per-residue model Mpipi,^62^ which has been validated to capture the phase behavior of disordered proteins. Based on these results, we trained a neural network classifier to predict whether a peptide phase separates. The sequence features used as neural network inputs were based on composition and spatial distribution of charged, aromatic, and strongly self-interacting residues (see Supplementary Information),^25^ and were obtained by a linear transformation from the one-hot encoded amino acid sequence. We then leveraged this predictor to filter new random peptides, retaining 200 sequences predicted to phase separate for the more computationally expensive simulations (Figure 1B). In these longer slab simulations (Figure 1C), we quantified *c*_dense_ and *γ* (evaluated from the computed pressure tensor^63^) for cases in which a dense phase formed. Based on this additional information, we updated the phase separation classifier and trained a regressor to predict *c*_dense_ and *γ*.

As a next step, we built on a previously developed method to identify globally optimal peptide sequences given the trained neural networks.^25^ The piecewise linear nature of the ReLU activation function allowed us to formulate the optimization as a mixed-integer linear programming (MILP) problem using the big-M reformulation.^64,65^ The regressor maps each sequence to its corresponding objective values, while the classifier is incorporated as a constraint to exclude non–phase separating sequences. This enables the identification of Pareto-optimal sequences, as predicted by the surrogate models, with mathematical guarantees of global optimality. Combined with an exploration strategy, this led to 100 new sequences to investigate in a new round of coarse-grained simulations. We stopped the optimization loop upon stagnation of the hypervolume of the Pareto front. More details regarding the computational pipeline can be found in Materials and Methods.

### Net charge can but does not necessarily break the phase separation–interfacial tension correlation

For the initial set of random peptides, *γ* showed a strong correlation with phase separation propensity (*c*_dense_), as expected. This correlation was also evident for all homopolymers that phase separate according to the Mpipi force field, ^62^ with *γ* appearing to increase nearly exponentially with *c*_dense_ (Figure 2A). Over the course of the optimization, peptides that strongly broke this correlation were identified (Figure 2B), and the hypervolume of the Pareto front converged after 8 iterations (Figure S1).

**Figure 2:**
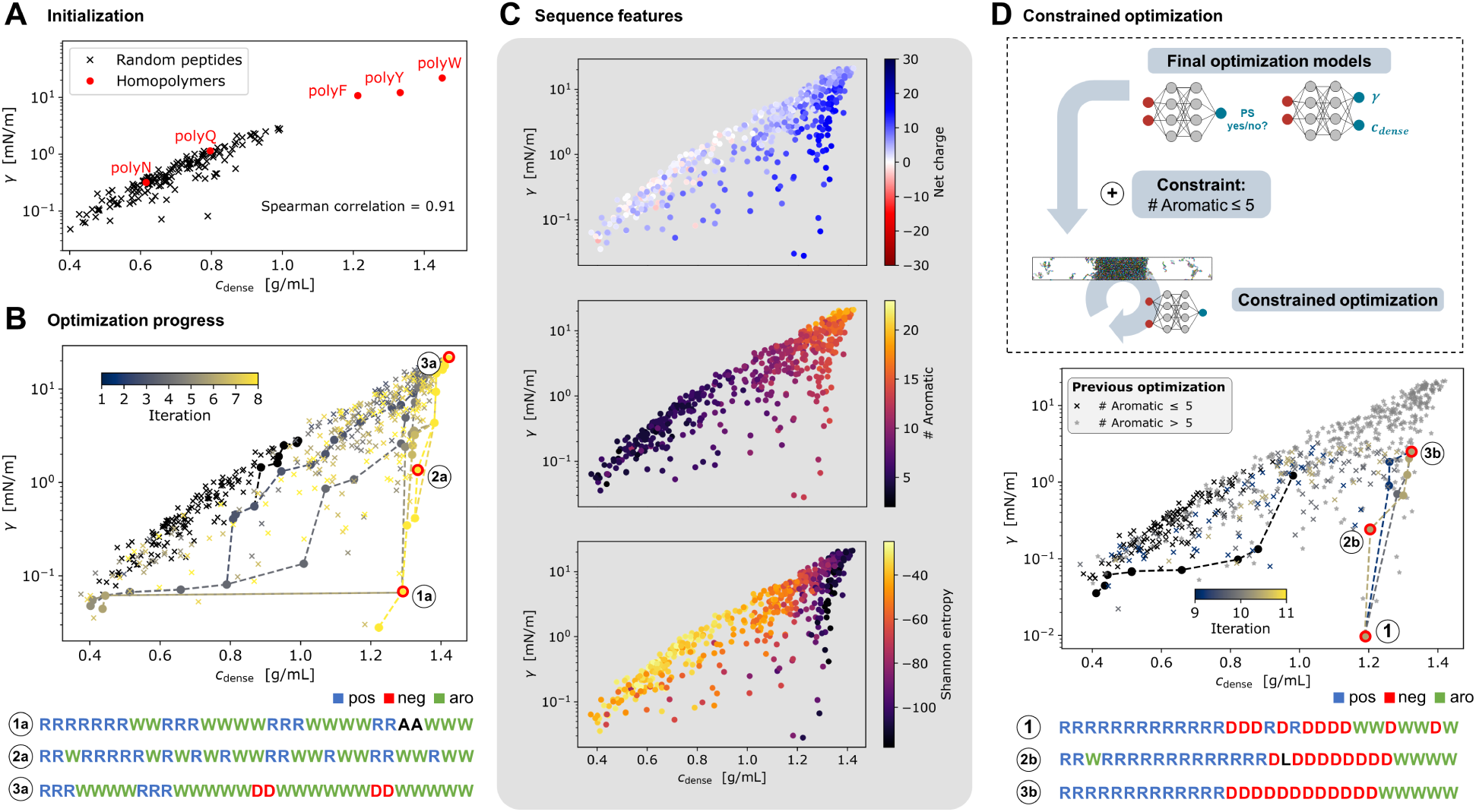
Optimization results for designing peptides with high phase separation propensity (*c*_dense_) and low interfacial tension (*γ*). (A) For random peptides and phase separating homopolymers, *γ* and *c*_dense_ are strongly correlated. (B) Optimization converges within 8 iterations, yielding arginineand tryptophan-rich Pareto-optimal sequences containing polyR patches. (C) Sequence feature analysis shows that net charge can, but does not necessarily, disrupt the *γ*–*c*_dense_ correlation. (D) Constrained optimization limiting the number of aromatic amino acids and incorporating prior data converges rapidly to similar arginine-rich, positively charged sequences.

Pareto-optimal peptides were predominantly composed of tryptophan (W) and arginine (R) residues, often adopting surfactant-like architectures with long arginine-rich segments (Figure 2B, Table S1). According to Mpipi, tryptophan exhibits strong interactions with arginine as well as the strongest self-interactions, resulting in a high phase separation propensity. Furthermore, studies have shown that clustering stickers within a sequence further enhances interaction strength,^66^ which is consistent with the presence of tryptophan patches observed here. More detailed analysis of sequence features reveals that all peptides breaking the *γ*–*c*_dense_ correlation exhibit a strong (mostly positive) net charge, primarily due to the presence of arginine residues (Figure 2C). Nevertheless, net charge alone is not sufficient to break this correlation, as similar net charges can be observed across three orders of magnitude for *γ*. It is also evident that a higher number of aromatic residues leads to an increase in *c*_dense_, and that the final Pareto-optimal sequences are composed of only a subset of amino acids, as reflected by their low Shannon entropy (Figure 2C).

Although the absence of aggregation flags by AGGRESCAN^61^ was imposed as a constraint during optimization, and a large fraction of sequences also passed the Waltz^59^ and TANGO^60^ filters, the high aromatic content of the peptides still raised concerns regarding aggregation. We therefore introduced an additional constraint, limiting sequences to a maximum of five aromatic residues. Leveraging the already trained model to guide this constrained optimization resulted in rapid convergence (Figure 2D), highlighting the effectiveness of our approach. The resulting Pareto-optimal sequences shared similar properties with the peptides from the first optimization, such as a positive net charge and the presence of aromatic and arginine patches. In addition, many sequences incorporated aspartic acid (D) residues, leading to more moderate, though still positive, net charges. Despite being limited to only five aromatic amino acids per sequence, the coexistence simulations indicate that these peptides can still form very dense phases with minimal interfacial tension. All Pareto-optimal sequences, including those obtained after applying the Waltz and TANGO filters, are provided in the Supplementary Information (Figure S2, Tables S1–S6).

For the next steps, we selected sequences 1 and 2b shown in Figure 2D.

### Sequence patterning can drastically influence interfacial tension

Since net charge appears necessary but not always sufficient for generating peptides with high phase separation propensity and minimal interfacial tension, we next investigated the effect of sequence patterning. In the following, we describe the results obtained with sequence 1 shown in Figure 2D, but similar findings were observed also with the sequence 2b (results shown in Figures S4-5). To isolate sequence patterning effects, we fixed the peptide composition in the next optimization step by adding a set of equality constraints to the optimization. Furthermore, in contrast to the previous setup, we reversed one optimization objective, switching from minimizing to maximizing *γ* (Figure 3A). With this strategy, we aimed to test whether it is possible to generate sequences that behave drastically differently from the original peptide, despite having identical composition.

**Figure 3:**
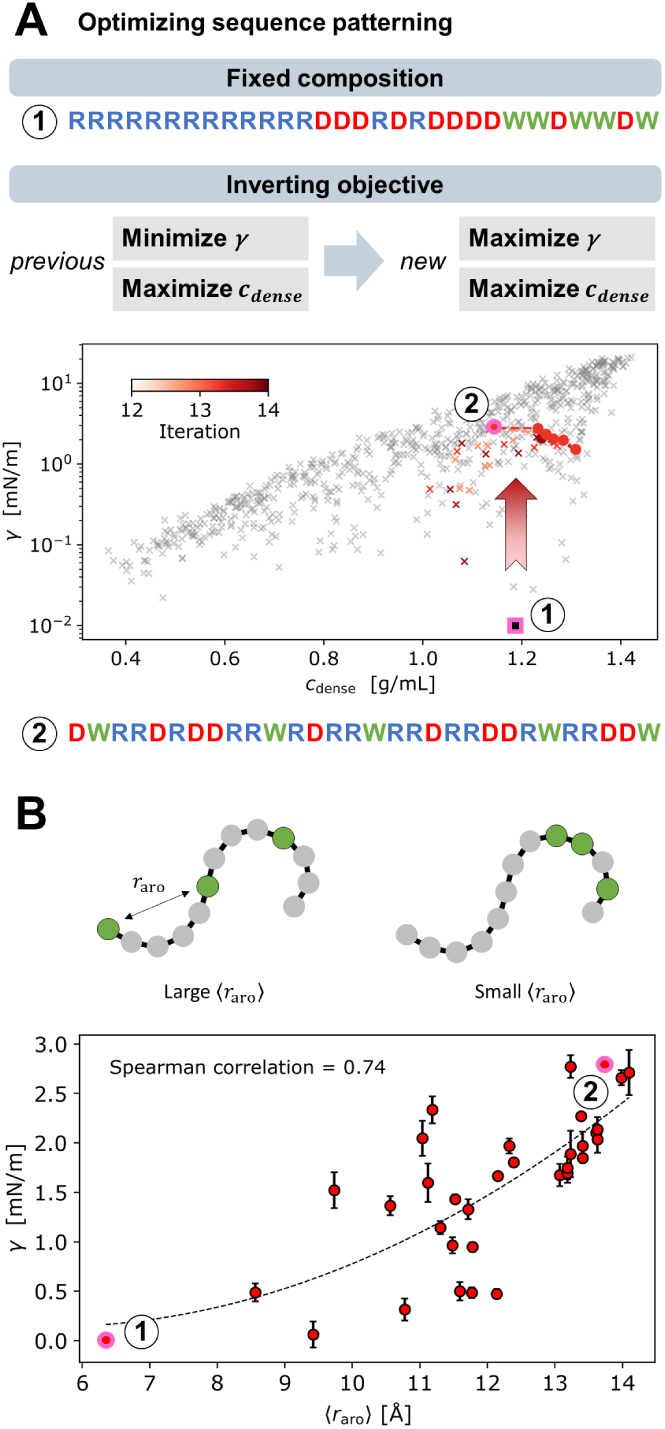
Varying interfacial tension with constant peptide composition. (A) In this third optimization, variation in sequence patterning was able to shift *γ* from low to high values. (B) The mean separation distance between aromatic residues ⟨*r*_aro_⟩ correlates with *γ*, indicating that clustering of tryptophans, and the corresponding long aromatic-free segments, enables low *γ* values.

As before, we built upon the previously available data and trained models to guide this next phase of optimization. The optimization process converged rapidly once again, resulting in a significant increase in *γ* through sequence modification. One example is the new Pareto-optimal sequence 2 shown in Figure 3A, which yields a much higher *γ* than the original version, while *c*_dense_ remains nearly constant. This particular sequence more closely resembles a random copolymer and exhibits reduced blockiness compared to the original peptide.

To more systematically examine which sequence patterning features determine interfacial tension, for all peptides we evaluated common descriptors related to patterning of charged or strongly self-interacting residues. Interestingly, neither sequence charge decoration (SCD),^67^ sequence hydropathy decoration (SHD),^68^ nor the charge blockiness parameter (*κ*)^69^ exhibited a significant correlation with interfacial tension (Figure S5). By contrast, analysis of surrogate model performance indicates that *γ* can be predicted with reasonable accuracy (Figure S3). This suggests that while the trained model captures the underlying physics of the problem, the observed behavior cannot be explained by these individual patterning features alone. We constructed a new simple feature that reasonably correlates with *γ*, namely the mean separation distance between two aromatic residues ⟨*r*_aro_⟩, estimated under the assumption of an ideal polymer chain as a zero-order approximation of inter-residue distances (Figure 3B, Supplementary Information). This feature suggests that low interfacial tension can be achieved when aromatic residues (here tryptophans) are clustered together, also allowing for the formation of longer segments devoid of these strong stickers. A more even distribution of aromatic residues along the sequence, thereby reducing sequence blockiness, generally results in high values for ⟨*r*_aro_⟩ and *γ*, as observed for sequence 2.

### Nanocondensate metastability depends on sequence patterning

Up to this point, all peptides have been investigated within slab coexistence simulations only, quantifying *c*_dense_ and *γ* without explicitly representing nanocondensates. As a next step, we investigated how these properties affect the evolution of the size distribution in a phase separating system. We proceed with sequences 1 and 2 shown in Figure 3 to analyze how different sequence patterning, with identical composition, influences nanocondensate formation *in silico*. For this purpose, we simulated the evolution of the size distribution by evenly distributing 1000 peptides in a cubic simulation box and allowing the system to evolve. At the beginning of the simulation, systems with peptide 1 and peptide 2 behaved similarly: oligomers rapidly formed, as reflected by the initial sharp decrease in the number of clusters (Figure 4A). At larger simulation times, the behavior of the two systems diverged: peptide 2 transitioned into one large droplet after ∼250 ns, while peptide 1 still showed six condensates at this point. This system remained in a state with three condensates during the last 500 ns of the simulation. The average size of peptide 2 condensates followed the scaling law ⟨*d*⟩ ∝ *t*^1^*^/^*^3^, expected from Ostwald ripening and Brownian-motion induced coalescence,^70^ whereas this behavior was not observed for peptide 1 (Figure 4A).

**Figure 4:**
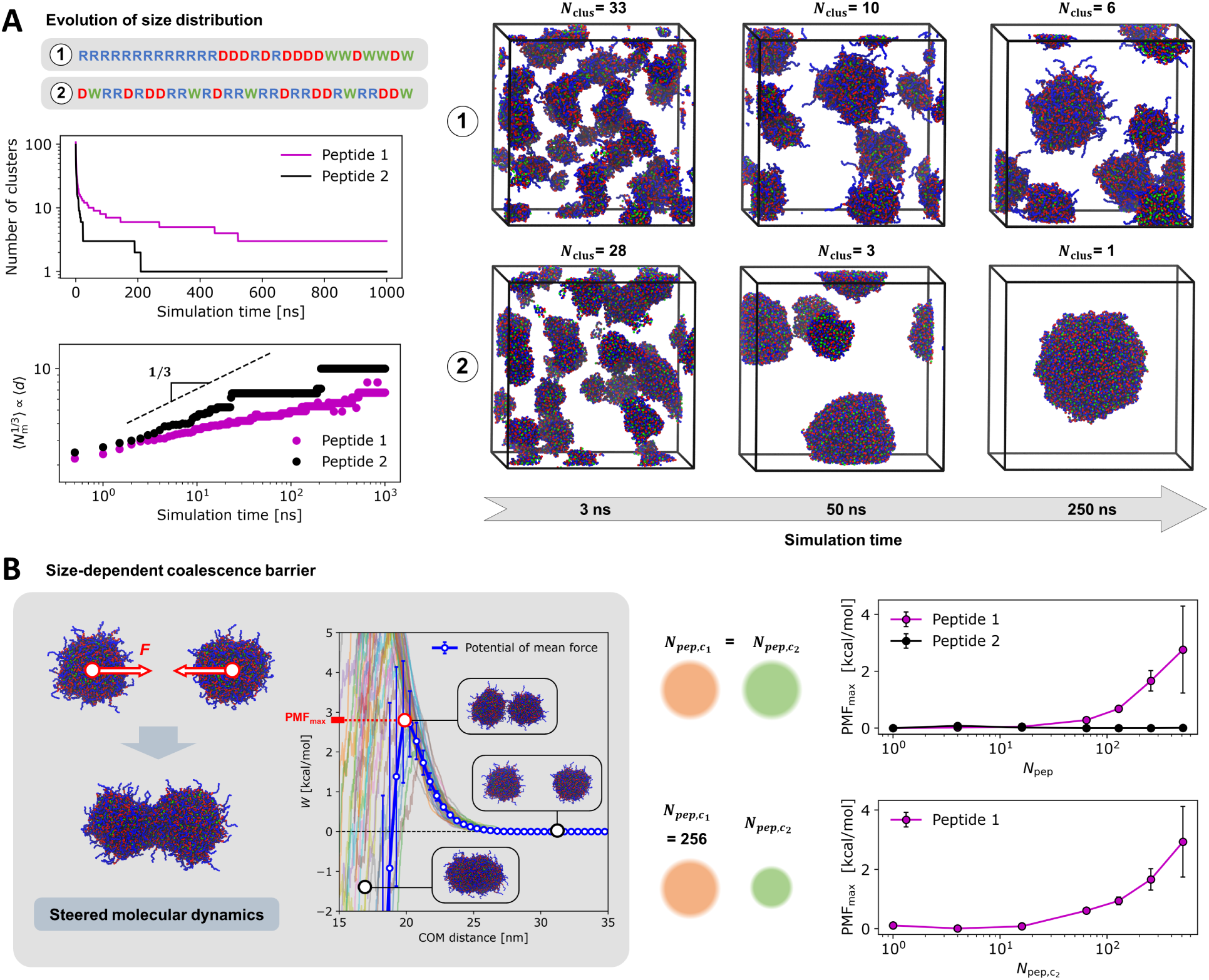
Analysis of nanocondensate metastability for low-*γ* peptide 1 and high-*γ* peptide 2, which share the same composition. (A) Time evolution of the condensate size distribution: the system with peptide 2 rapidly coalesces into a single dense phase, whereas the system with peptide 1 becomes trapped in a state with multiple smaller nanocondensates. (B) Coalescence barriers from steered molecular dynamics: for peptide 1, a barrier exists for the coalescence of two large nanocondensates; however, this barrier is absent during events involving the interaction of a large nanocondensate with a small nanocondensate or between two small nanocondensates. No barrier is detected for peptide 2.

The number of free monomers remained negligible in both cases, indicating absence of Ostwald ripening at the simulated timescale and at these high total concentrations. The distinct evolution of the size distribution is therefore governed by coalescence, which appears to be inhibited for the low-*γ* peptide 1. We analyzed this difference in more detail by quantifying a potential coalescence barrier as a function of condensate size for both peptides. This was achieved by repeatedly simulating coalescence events between two condensates using steered molecular dynamics, applying the center-of-mass (COM) distance as the collective variable (CV). We constructed the potential of mean force (PMF) by quantifying the work performed during this nonequilibrium process and subsequently applying Jarzynski’s equality.^71,72^ We selected this method since we were not interested in the full free energy profile across the entire CV range, but only in the PMF maximum, which we used as a measure of the coalescence barrier (Figure 4B).

When varying the size of the coalescing nanocondensates, it becomes apparent that for peptide 1 a potential barrier emerges at large condensate sizes. For peptide 2 this barrier is absent, even though its composition, and thus its net charge, is identical to peptide 1. In the next step, we fixed the size of one condensate at 256 peptides and varied only the size of the second condensate (Figure 4B). We find that for peptide 1 the coalescence barrier exists only for large-large coalescence events, whereas monomers and smaller oligomers can still merge readily with large condensates. Overall, these findings explain the distinct behavior of peptide 1 and 2 shown in Figure 4A and suggest that peptide 1 forms metastable nanocondensates in solution, because Ostwald ripening is slow and coalescence is inhibited once nanocondensates exceed a size threshold. Moreover, the free energy minimum corresponded always to the coalesced state, consistent with thermodynamic equilibrium favoring a macroscopic dense phase (Figure S6-8). The observed nanocondensates of peptide 1 therefore clearly exist as metastable, kinetically trapped species.

### Electrostatics-driven interfacial structuring depends on condensate size and sequence patterning

The results described above show that the low-*γ* peptide 1 exhibits a size-dependent coalescence barrier that leads to metastable nanocondensates, whereas the high-*γ* peptide 2 does not, despite having the same composition. Next, we aimed at understanding the molecular basis behind this difference by simulating nanocondensates of different sizes and analyzing their structure. For peptide 1, we observed that the organization of molecules at the interface changes as a function of nanocondensate size (Figure 5A): in particular, the poly-arginine (R) tails are progressively more excluded with increasing the size of condensates. We illustrate this change by plotting the density of different residue types against the center-of-mass (COM) distance in a condensate consisting of 1024 peptides (Figure 5A). The outer interface layer is mainly composed of arginines, while the tryptophan (W) residues are more buried within the condensate. For peptide 2, by contrast, the composition is roughly equal at all distances (Figure 5A, Figure S9-10).

**Figure 5:**
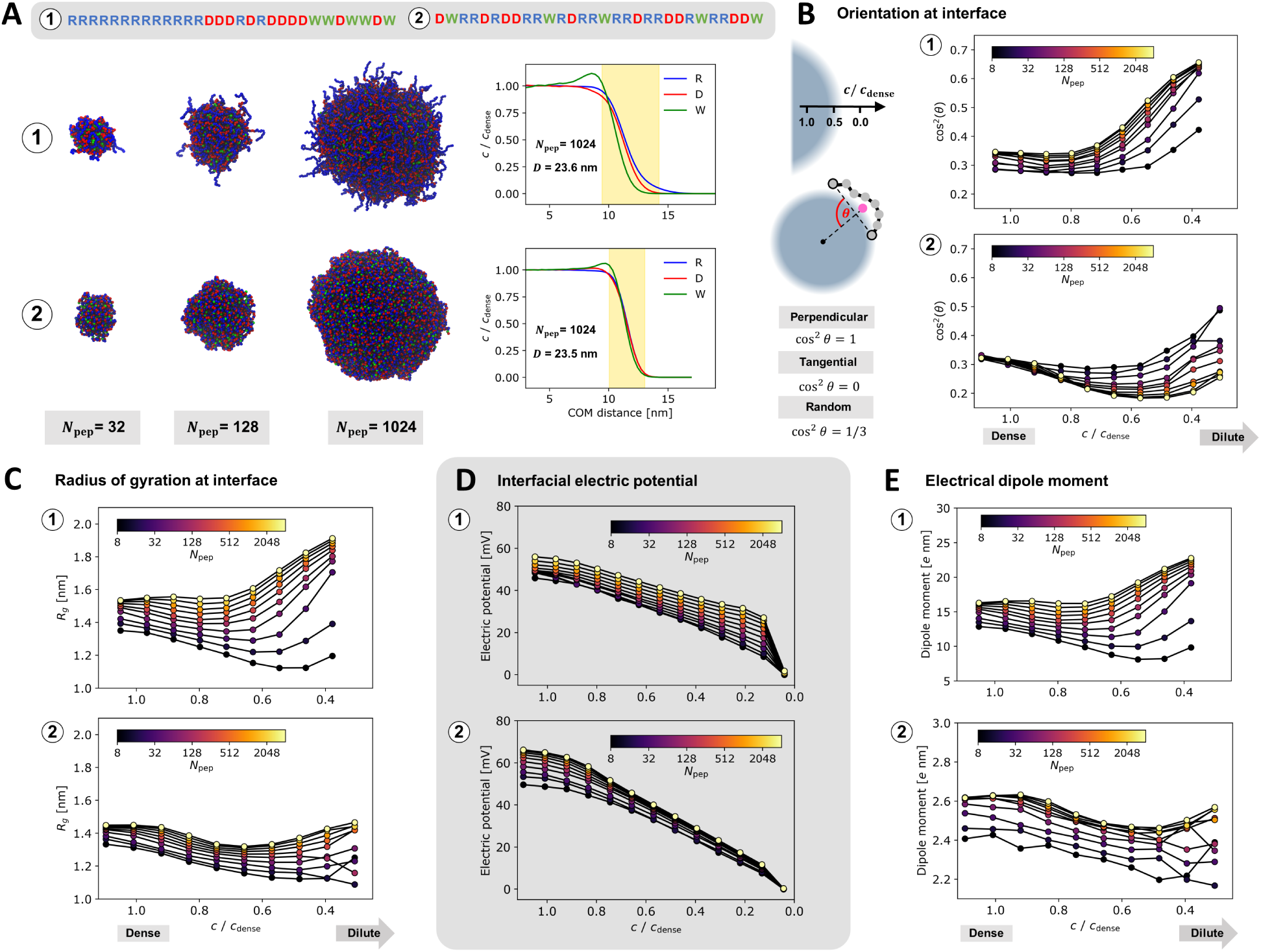
Investigation of size-dependent interfacial structuring. (A) Simulations of individual nanocondensates of varying sizes (represented by the number of peptide molecules *N*_pep_) reveal that arginine is excluded from the dense phase of peptide 1, especially in condensates with larger sizes. (B) The orientation of interfacial molecules changes with increasing condensate size: peptide 1 becomes more perpendicular, while peptide 2 more tangential. (C) The molecular expansion at the interface is also size-dependent, with peptide 1 becoming increasingly more expanded at the interface compared to bulk with increasing the size of the condensate. (D) The interfacial electric potential arising from the peptide’s net charge (+5) drives exclusion of arginine residues. (E) Electric dipole moment shows a trend similar to the radius of gyration. Values are generally much higher for peptide 1 than for peptide 2.

To get further molecular insights, we next analyzed the molecular orientation as a function of the position within the condensate considering different condensate sizes (Figure 5B). Building on the approach of Farag et al.,^73^ we quantified the angle *θ* defined by the vector connecting the two peptide ends and the vector between the condensate COM and the peptide COM. We did not calculate *θ* simply as a function of the distance between peptide and condensate COM because individual nanocondensate conformations are not necessarily radially symmetric because of nanocondensate shape fluctuations. Therefore, averaging properties based simply on COM distance can introduce artifacts, such as a widening of the interfacial region in larger droplets, and compromise comparison across distinct condensate sizes. We instead defined spatial localization based on the smoothed local density at the peptide COM position, relative to the dense phase (see also Materials and Methods). A value of ∼1.0 corresponds to the dense phase and ∼0.0 to the dilute phase, with the interfacial region lying in between. For both peptides, chains in the dense phase adopt a nearly random orientation with cos^2^(*θ*) ≈ 1*/*3. At the interface, the orientation clearly depends on condensate size and converges for larger condensates in both cases. In the case of peptide 1, molecules in large condensates are more perpendicular to the condensate surface, consistent with the arginine residues being excluded from the dense phase. By contrast, for peptide 2 the interfacial molecules become increasingly aligned tangentially. This emergent interfacial structuring can also be detected by considering the peptide radius of gyration, which also depends on condensate size (Figure 5C): for peptide 1, chains are increasingly expanded at the interface compared to the bulk in condensates exceeding 32 peptide copies. In contrast, for peptide 2, chains at the interface are similar to or more compact than those in the dense phase.

The fact that poly-arginine is excluded from the dense phase for peptide 1, accompanied by molecular alignment and expansion, indicates that electrostatics plays a key role in interfacial structuring. Indeed, the condensate-forming peptide is positively charged (net charge of +5), leading to long-range electrostatic repulsion of arginine from the condensate. To get further molecular insights, we quantified the interfacial electric potential, which has been suggested to play key roles in a variety of processes such as promotion of redox reactions^38,46^ (Figure 5D). The potential inside the condensate is positive, driving arginine residues towards lower and aspartic acid residues towards higher potential. We observed that, at a fixed position relative to the interface, the electric potential increases with condensate size for both peptides. A contributing factor for this size dependence is the long-range character of electrostatic interactions together with the change of surface curvature, ^74^ leading to higher potentials with increasing condensate size. This potential and the corresponding electric field can lead to the alignment of molecules,^75^ explaining the observed behavior (Figure 5C). Consistent with the trends of the electrical field and radius of gyration, peptide 1 also exhibits a higher electric dipole moment at the interface (Figure 5E).

In summary, our results show that the condensate size–dependent interfacial electric potential and resulting field lead to emergent interfacial structuring. The sequence blockiness of peptide 1 and consequent polarizability cause molecular alignment, expansion, and exclusion of arginine segments from the dense phase. This structuring rationalizes the low interfacial tension observed in the large-condensate limit (slab simulations), as well as the size-dependent coalescence barrier arising from electrostatic repulsion of exposed arginine residues. The structuring is fundamentally different for peptide 2 condensates, where strong stickers (tryptophans) are exposed at the surface, leading to high *γ* and the absence of any significant coalescence barrier.

The insights of our work share analogies with findings observed with natural proteins that have been shown to form nanocondensates, demonstrating the possible generalizability of these mechanisms. An example is *α*-synuclein, which forms nanocondensates in subsaturated solutions.^8^ Similar to our designed peptides, the protein possesses a negatively charged tail (C-terminus) which matches the net negative charge of the protein. Mutation of this tail and decrease of its charge promotes phase separation over nanocondensate formation, suggesting that a similar mechanism stabilizes the nanocondensate state. Another example where mutations had a decoupled effect on nanocondensate formation and phase separation is FUS:^10^ increasing the charge of the C-terminus, which also matches the net charge of the protein, favors nanocondensates over phase separation. Here, we investigated the WT and the proposed mutant^10^ of FUS in more detail, and observed that the mutation in fact reduces interfacial tension and also leads to distinct interfacial structuring for larger nanocondensates (Figure S11). This indicates that net charge, in combination with sequence segments matching the scaffold’s net charge, is the mechanism behind nanocondensate metastability not only for our designed peptides but likely also for natural protein systems.

### Experimental validation of the designed peptides

To validate the computational predictions, we experimentally characterized the low-*γ* peptide 1 and high-*γ* peptide 2. We began by investigating their phase separation propensity using bright-field microscopy. Both peptides underwent macroscopic phase separation in a concentration-dependent manner, which was also sensitive to ionic strength. A minimum ionic strength of 150 mM and 350 mM was required for phase separation of peptides 1 and 2, respectively, within the tested peptide concentration range (Figure 6A, Figure S12-13).

**Figure 6:**
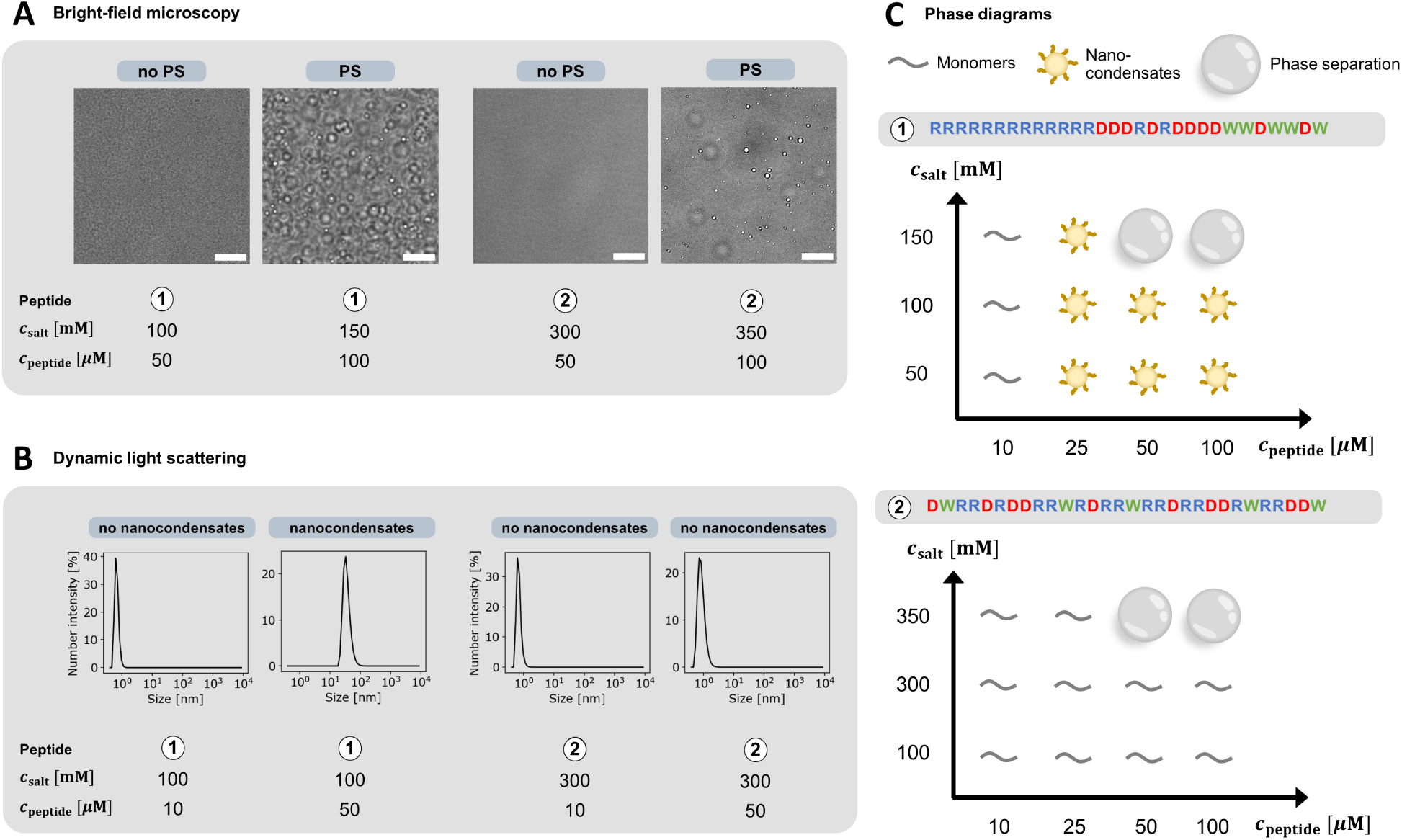
Experimental validation of the effect of sequence patterning on nanocondensate formation. (A) Representative bright-field microscopy images reporting on macroscopic phase separation behavior. Scale bars: 10 *µ*m. (B) Representative size distributions as measured by dynamic light scattering (DLS). Peptide 1 forms nanocondensates under conditions where macroscopic phase separation is absent, whereas peptide 2 does not. (C) Schematic phase diagram summarizing the results obtained with bright-field microscopy and DLS for both peptides. Peptide 1 exhibits a broad nanocondensate region, which is absent for peptide 2, in agreement with computational predictions.

The higher phase separation propensity of peptide 1 compared to peptide 2, with phase separation occurring at a lower ionic strength at the same peptide concentration, qualitatively aligns with the computational predictions. Indeed, in the slab coexistence simulations, *c*_dense_ was slightly higher for peptide 1 than for peptide 2 (Figure 3A).

We then quantified the size distribution under conditions where no micron-scale condensates were visible, using dynamic light scattering (DLS) (Figure 6B, Figure S14-15). We found that peptide 1 forms nanocondensates of approximately 30 nm across a broad range of conditions where no macroscopic phase separation is observed. The peptide remains within the monomeric size range (1–2 nm) only at the lowest overall concentration tested.

In contrast, peptide 2 transitions directly from a solution of monomers to macroscopic phase separation, without forming any mesoscale cluster species (Figure 6A-C). These experiments confirm our predictions that only peptide 1 is capable of forming metastable nanocondensates, despite both peptides sharing the same composition. Additionally, for peptide 1, we observed a time-dependent shift in the size distribution (Figure S16), consistent with reports for other protein nanocondensate systems,^8–10^ confirming the metastable character of this species.

Overall, the agreement between computational predictions and experiments confirms our design approach as well as the proposed mechanism underlying nanocondensate formation. While peptide 1 shows a metastable nanocondensate region within its phase diagram, peptide 2 with the same composition exhibits a direct transition from the monomeric size range to macroscopic phase separation.

## Discussion

In this study, we developed and applied a computational pipeline leveraging high-throughput coarse-grained simulations, machine learning, and mixed-integer linear programming to design peptides that form metastable nanocondensates under a broad range of operating conditions. To this aim, we generated and experimentally validated peptides with high phase separation propensity (high *c*_dense_) but low interfacial tension (*γ*), breaking the correlation which is typically observed among these two properties. This leads to a strong driving force to form nanocondensates, without transitioning into a macroscopic dense phase.

Our findings indicate that the net charge of the condensate-forming peptide combined with sequence blockiness are effective drivers for breaking the *c*_dense_-*γ* correlation and for inducing the formation of metastable nanocondensates instead of a macroscopic dense phase. We also demonstrate that merely changing sequence patterning can determine whether these assemblies form (Figure 7).

**Figure 7:**
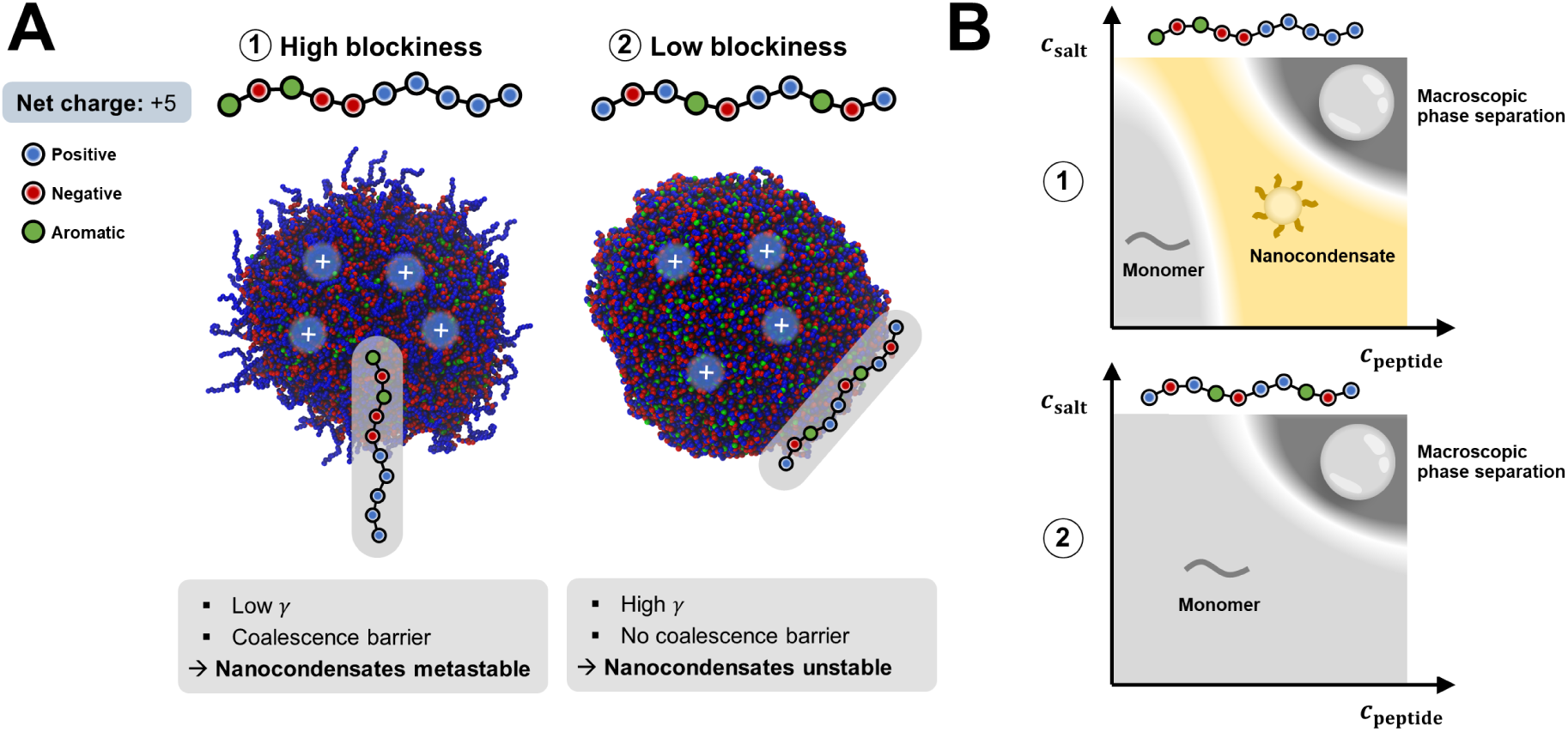
Summary of the effect of sequence patterning on nanocondensate formation. (A) Molecular mechanism: The high blockiness of peptide 1 leads to molecular alignment and exclusion of the positively charged segment matching the dense-phase charge. This leads to low interfacial tension *γ*, directly slowing Ostwald ripening, and inhibited coalescence. In contrast, the architecture of peptide 2 promotes tangential alignment with all residue types exposed at the surface, yielding fundamentally different behavior. (B) Phase behavior: in contrast to peptide 2, peptide 1 shows a broad region with metastable nanocondensates.

Our analysis has further highlighted the molecular basis behind the formation of these metastable nanocondensates. The interfacial electric potential, together with sequence segments whose charge matches the nanocondensate’s net charge, leads to a condensate size-dependent interfacial structuring that excludes these segments. This emergent molecular alignment is at the origin of the low *γ* values and the inhibition of Ostwald ripening. It also hinders the coalescence of condensates beyond a threshold size through electrostatic repulsion of the excluded charged segments. Molecular alignment and expansion at condensate interfaces have previously been observed *in silico*.^73,76^ Here, we demonstrate that this behavior strongly depends on sequence patterning and condensate size. This observed size-dependent interfacial structuring indicates that both kinetic and thermodynamic surface properties vary with condensate size. This directly violates a central assumption of classical nucleation theory, namely that interfacial tension is size-independent, and helps rationalize why nucleation behaviour in condensates can deviate markedly from classical predictions.^77^ Importantly, we have demonstrated that our mechanism applies not only to the designed peptides but also to natural nanocondensate-forming proteins, implying its potential generalizability. The broad applicability of this mechanism is also supported by its alignment with key generic features of condensate systems, such as the extremely low interfacial tension of condensates formed by natural proteins.^78^ Moreover, experiments have shown that nanocondensates can exchange molecules via the dilute phase. ^10^ This is fully compatible with our mechanism: the coalescence barrier suppresses fusion between large nanocondensates while still allowing single-monomer exchange, and Ostwald ripening remains slow due to the low interfacial tension *γ*.

Our proposed mechanism converges with prior conceptual frameworks that have been used to explain nanocondensate formation. Our study showed that homopolymers could not break the *c*_dense_-*γ* correlation, thus implying that different interaction sites characterized by distinct energy scales are required for nanocondensate formation, as previously suggested.^10^ Moreover, our analysis has revealed that the low interfacial tension arises from distinct conformations of molecules at the interface, as discussed previously.^14,21^ Furthermore, consistent with our findings, the net charge of condensates has also been proposed as a key determinant favoring the formation of nanocondensates rather than macroscopic condensates.^19,20,29^ Finally, saturation of valences and the lack of exposed strong stickers at the interface have been suggested as sources of kinetic arrest,^30^ consistent with the behavior of our designed peptide 1. Therefore, our study integrates these concepts and grounds them in concrete examples, thereby deepening our understanding of the mechanisms underlying nanocondensate formation.

## Conclusions

Using a newly developed computational pipeline, we designed peptides that form metastable nanocondensates across a broad range of operating conditions. This design process revealed that nanocondensate metastability is governed by net charge and sequence blockiness. We unraveled the molecular mechanism underlying the observed nanocondensate metastability, showing the presence of an electrostatics-driven, size-dependent interfacial structuring that leads to inhibited ripening and coalescence. This mechanism not only unifies previously proposed explanations of condensate stability but is also consistent with key observations in many other protein nanocondensate systems.

Beyond advancing our understanding of nanocondensate systems, the findings and methodology presented here enable the rational design of nanocondensates in the nanometer size range observed in cells.^18^ We envision that the design of metastable nanocondensates with tunable size distributions offer promising opportunities in bioengineering and biotechnology, for instance in drug delivery and optimization of biochemical reactions.^34^

## Materials and Methods

### Molecular simulations

Molecular simulations were carried out using the one-bead-per-residue force field Mpipi^62^ in an NVT ensemble, applying a Langevin thermostat^79^ at 300 K with a relaxation time of 100 ps and timestep of 10 fs. Simulations were run using the LAMMPS Molecular Dynamics package (version 2nd of August 2023)^80,81^ and trajectory analysis was performed using Python (version 3.11) and the MDanalysis library (version 2.9.0).^82,83^

#### Slab simulations

Due to the variety of peptide sequences, a robust setup for the slab coexistence simulation was crucial: A total of 325 peptide copies were placed in a periodic box with initial dimensions 113×113×290 Å^3^, which was then elongated in the *z* direction to final dimensions of 113×113×890 Å^3^. The peptide slab was compressed by applying harmonic walls along the *z* direction (force constant: 0.1 kcal mol*^−^*^1^ Å*^−^*^2^, width: 150 Å), followed by energy minimization and a short equilibration of 3 ns. Harmonic walls were then extended to a width of 250 Å, and equilibrated for another 5 ns, after which they were removed. For the unrestrained system, a final equilibration of 20 ns was performed, followed by an 80 ns production run. The binned density along the *z* direction and the pressure tensor were collected. Phase separation was detected when the density in the center of the simulation box significantly exceeded the average density, as determined by a one-sided t-test. Especially for low-*γ* systems, fracture and subsequent fusion of the slab can occur. Frames with a fractured dense phase were identified by the presence of multiple peaks in the density profile using find peaks implemented in SciPy,^84^ and discarded for further analysis. Cases in which the dense phase was fractured for more than 50 % of the trajectory were classified as no phase separation. In cases of phase separation, the symmetric density profile was mirrored and averaged to yield a single half-profile, which was then fit with a sigmoid function, with the high plateau corresponding to *c*_dense_. Interfacial tension was determined from the pressure tensor as described by Tejedor et al.^63^ Simulations yielding negative interfacial tensions, indicating a kinetically arrested but unstable slab, were discarded before training the surrogate model in the next step. For each peptide, three simulations with different seeds were run. For the simulations assessing only phase separation (Figure 1B), no compression with harmonic walls was applied, the production run was limited to 20 ns, and each simulation was performed only once.

#### Time evolution of size distribution

A total of 1000 peptide molecules were evenly distributed in a periodic cubic simulation box with an edge length of 419 Å. Energy minimization was followed directly by a 1 *µ*s production simulation. The number of clusters in each frame, as well as the number of peptides per cluster, were quantified using the DBSCAN^85^ algorithm with peptide COM coordinates as input and a distance cutoff of 30 Å, implemented in scikit-learn (version 1.6.1),^86^ followed by smoothing across multiple frames.

#### Interfacial structure analysis

Nanocondensates of different sizes were prepared by randomly placing monomers in a cubic box and applying a linear potential, and thus a constant force, on the radius of gyration of all molecules, leading to all peptides being in one condensate. The bias potential was then removed, followed by 50 ns of equilibration and a 250 ns production run. As described in the main text, orientation, radius of gyration, and electrical dipole moment were calculated as a function of local density at the COM position for each peptide in the condensate. For this purpose, the density field was calculated for each frame by constructing a grid with 20 Å spacing, followed by smoothing with a second-order spline filter. The density at each peptide COM was then obtained by second-order spline interpolation. Electrical dipole moment **p** was calculated using:

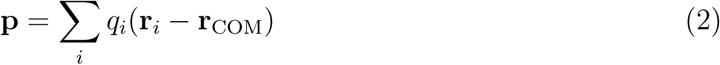

Electric potential was quantified by randomly inserting 200k unit-charge probes into the simulation box, followed by calculating their interaction energies using a lower distance cutoff of 5 Å to avoid particle collisions, inspired by Tsanai et al.^87^

#### Quantifying coalescence barrier

For quantification of the potential barrier for coalescence, two nanocondensates prepared following the same protocol described in the previous section were placed in a simulation box. A steered molecular dynamics simulation was then performed using a moving harmonic restraint applied to their center-of-mass separation distance. The force constant was set to *k* = *N*_pep_ × 5 kcal mol*^−^*^1^ Å*^−^*^2^, where *N*_pep_ is the number of peptides per condensate, and the pulling velocity was set to 2 Å ns*^−^*^1^. In contrast to previous simulations, a Nose-Hoover thermostat (300 K, 5 ps) was applied to improve convergence, removing the Langevin friction term. For condensate sizes *<*128 monomers, 100 simulations with different random seeds were carried out. For larger systems, 50 simulations were performed to limit computational cost. The quantified work *W* from each run was transformed into the potential of mean force (PMF) Ψ using Jarzynski’s equality,^71^ more specifically its second-order cumulant expansion, which provides more accurate estimates than the exponential average when only a limited number of trajectories is available:^72^

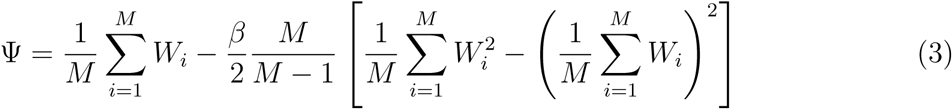

where *M* is the number of sampled trajectories, standard deviations are estimated through error propagation. In Figures S6–8, a few outlier trajectories with high work values are observed, originating from stochastic dissociation of the assembly. The influence of these outliers is marginal and was therefore neglected, except for clusters of size 4, where dissociation events occurred frequently within the simulation time. In this case, dissociation was prevented by restraining the radius of gyration to *<*50 Å using harmonic walls.

### Machine learning and optimization

The machine-learning and optimization setup builds on previous work, where multi-output neural networks were used in conjunction with mixed-integer linear programming (MILP) to identify globally optimal peptide sequences.^25^ We use the same set of 44 engineered features (Supplementary Information) to train two different surrogate models on all available data at each iteration: The classifier predicting whether a peptide phase separates is trained on all peptides and consists of two hidden layers with 10 and 5 neurons respectively, followed by a sigmoid layer that converts the final-layer logit into a probability estimate. The regressor, which predicts *c*_dense_ and ln *γ*, is trained only on the phase-separating peptides and consists of two hidden layers with 30 neurons each. Both models were trained with a batch size of 32 and the Adam optimizer,^88^ using PyTorch^89^ (version 2.3.1) and scikit-learn^86^ (version 1.5.1). A multi-step learning rate scheduler was employed, decaying the learning rate by 0.9 at epochs 200, 400, 600, and 800. Features and regression objectives were normalized to the range [-1,1]. Smooth L1 loss was used for the regressor, and Binary Cross-Entropy loss for the classifier. 10% of the training data was randomly set aside to calculate validation loss, and the model with the lowest value was selected. Initial learning rate and weight decay were optimized using an 80/20 random train-test split, and final model performance evaluated on a separate holdout set (20%). For the optimization step, the models were trained on all data available up to this point. To mitigate potential numerical issues in the optimization part, all weights and biases with absolute values below 10*^−^*^3^ were set to zero.

For the MILP optimization, the trained neural networks were incorporated into the algebraic modeling framework Pyomo^90,91^ (version 6.7.3) using OMLT^65^ (version 1.1). For the exploitation component of the optimization, the Pareto front (50 sequences), based on the predicted *c*_dense_ and ln *γ* objectives, was constructed with Gurobi^92^ using the *ε*-constrained method, from which 25 well-spaced points were selected for the next iteration. For the exploration component, between 2 and 20 positions (chosen uniformly at random) were constrained to random amino acids, and the remaining positions were optimized. This optimization was performed for 150 sequences per iteration using the weighted-sum method with random weights, and a well-spaced subset of 75 sequences was selected. Two constraints were enforced during both these optimizations: first, the AGGRESCAN predictor was integrated to avoid generating aggregation-prone sequences ^25,61^ and second, the trained phase-separation classifier was used to exclude sequences that are not predicted to phase separate. More specifically, we required that the classifier’s predicted phase-separation probability exceeds 75%. This leads to a total of 100 peptide sequences for a new round of molecular dynamics simulations every iteration.

For the optimizations with fixed composition, we added additional equality constraints specifying the total number of residues of type X. Due to this highly reduced design space, we constructed the Pareto front using only 5 sequences and replaced the exploration strategy by generating 10 random shuffles at each iteration, for both peptide 1 and 2b. Because each iteration added only a marginal amount of new data relative to the existing dataset, the newly acquired data was resampled 10 times by adding Gaussian noise based on the mean and standard deviation of the simulation replicates.^93^ In general, the optimizations were stopped once the hypervolume stagnated.

### Dynamic light scattering

Peptides 1 and 2 were synthesized by Genscript (China) and dissolved in ultrapure water to give a final stock concentration of 1 mM. Samples were then diluted with filtered 20 mM Tris buffer at pH = 7 in 1.5 mL Eppendorf tubes to achieve desired peptide concentrations (ranging from 10 to 100 *µ*M). The target ionic strength (ranging from 50 to 350 mM) was adjusted by mixing suitable volumes of Tris buffer solutions with and without 5 M NaCl. The samples were subsequently equilibrated for 2 h at room temperature, and then centrifuged at room temperature for 30 min at 4000 rcf. The supernatant was then collected for analysis in disposable cuvettes (Brand GmbH) on a Zetasizer Nano-ZS (Malvern).

### Brightfield microscopy

20 *µ*L samples were prepared in a 384 MicroWell Glass Bottom MatriPlate (Azenta Life Sciences) at the desired buffer and peptide conditions following the same protocol used for dynamic light scattering measurements. After 2 h incubation at room temperature, during which the well plate was sealed, samples were imaged using brightfield microscopy (Eclipse Ti-E, Nikon) using a 60x oil objective (FI Plan Apo Lambda NA 1.4, Nikon).

## Data availability

Sequence feature definitions and analyses, machine learning model performance, optimization progress and Pareto-optimal peptide sequences, as well as molecular simulation, microscopy, and dynamic light scattering data are provided in the Supplementary Information.

## Acknowledgements

We kindly acknowledge the European Research Council through the Horizon 2020 research and innovation programme (grant agreement No. 101002094) for financial support. Computations were performed on the high-performance computing cluster Euler by ETH Zurich.

## Author contributions

T.N.S. and P.A. designed the conceptual framework of the study. T.N.S. and M.A.B. developed and applied the computational pipeline and performed the simulations. F.S., K.M., and M.R. performed the experiments. T.N.S., F.S., and P.A. wrote the manuscript with contributions from all authors. P.A. acquired funding and supervised the project.

