## Supplementary Information for "Engineering nanocondensate formation through sequence composition and patterning"

### 1 Sequence features

#### 1.1 Machine learning models

The 44 sequence features were adapted from previous work.[1] They encode composition as well as patterning of charged, aromatic, and strongly self-interacting residues. Importantly, they can all be derived by a linear transformation of the one-hot encoded amino acid sequence, which is crucial for the MILP optimization part.

- **20 features**: Number of amino acids of type X.

$$AA_X = \sum_{i=1}^{N} \delta_{a_i,X} \qquad \delta_{a_i,X} = \begin{cases} 1, & a_i = X \\ 0, & a_i \neq X \end{cases} \tag{1}$$

- **4 features**: Total number and distribution of positive charges along the sequence. $d_i$ is the distance from the center of the peptide, $d_i \in [-14.5, 14.5]$.

$$P_n = \sum_{i=1}^{N} \text{charge}_i^+ * d_i^n \qquad n \in [0, 3] \tag{2}$$

- **4 features**: Total number and distribution of negative charges.

$$N_n = \sum_{i=1}^{N} \text{charge}_i^- * d_i^n \qquad n \in [0, 3] \tag{3}$$

- **4 features**: Total number and distribution of aromatic residues.

$$A_n = \sum_{i=1}^{N} \text{aromatic}_i * d_i^n \qquad n \in [0, 3] \tag{4}$$

- **4 features**: Distribution of self-interacting residues. $\phi(r)$ is the Wang-Frenkel potential[2] with residue-specific parameters taken from Mpipi.[3]

$$E_n = \sum_{i=1}^{N} \epsilon_i * d_i^n \qquad \epsilon = \int_{\sigma}^{3\sigma} \phi(r)dr \qquad n \in [0,3] \tag{5}$$

- **4 features**: Total and distribution of molecular weight.

$$M_n = \sum_{i=1}^{N} MW_i * d_i^n \qquad n \in [0,3] \tag{6}$$

- **4 features**: Features derived from the commonly used SCD[4] and SHD.[5]

$$SHD = \sum_{i=1}^{N} \sum_{j=i+1}^{N} (\epsilon_i + \epsilon_j)(j-i)^{-1} \tag{7}$$

$$SPD = \sum_{i=1}^{N} \sum_{j=i+1}^{N} (\text{charge}_i^+ + \text{charge}_j^+)(j-i)^{-1} \tag{8}$$

$$SND = \sum_{i=1}^{N} \sum_{j=i+1}^{N} (\text{charge}_i^- + \text{charge}_j^-)(j-i)^{-1} \tag{9}$$

$$SPND = \sum_{i=1}^{N} \sum_{j \neq i}^{N} (\text{charge}_i^+ + \text{charge}_j^-)|j-i|^{-1} \tag{10}$$

#### 1.2 Sequence patterning analysis

For the analysis performed in Figure 3B and Figure S5, several existing features were used, which are defined as:[4–6]

$$SCD = \frac{1}{N} \sum_{i=1}^{N} \sum_{j=i+1}^{N} q_i q_j (j-i)^{1/2} \tag{11}$$

$$SHD = \frac{1}{N} \sum_{i=1}^{N} \sum_{j=i+1}^{N} (\epsilon_i + \epsilon_j)(j-i)^{-1} \tag{12}$$

$$\kappa = \left(\frac{\delta}{\delta_{\max}}\right) \qquad \text{with} \qquad \delta = \frac{\sum_{i=1}^{N_{\text{blob}}} (\sigma_i - \sigma)^2}{N_{\text{blob}}} \quad \sigma_i = \frac{(f_+ - f_-)_i^2}{(f_+ + f_-)_i^2} \tag{13}$$

with the standard value of 5 for the blob size $N_{\text{blob}}$. Finally, the mean aromatic separation distance $\langle r_{\text{aro}} \rangle$ was calculated assuming an ideal polymer chain which follows a 3-dimensional random walk:[7]

$$\langle r_{\text{aro}} \rangle = \frac{2}{N_{\text{aro}}(N_{\text{aro}} - 1)} \sum_{i=1}^{N_{\text{aro}}} \sum_{j=i+1}^{N_{\text{aro}}} b(j-i)^{1/2} \tag{14}$$

with a C$\alpha$ separation distance $b$ of 3.8 Å.

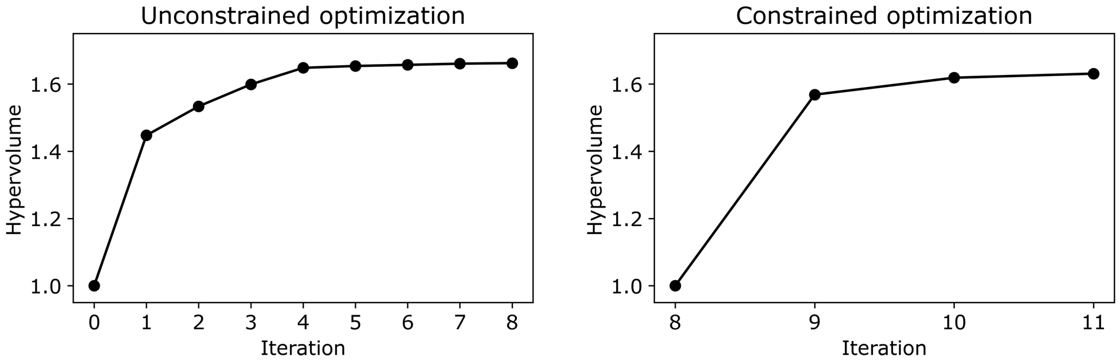


Fig. S1: Evolution of the normalized hypervolume for the optimizations shown in Figure 2.

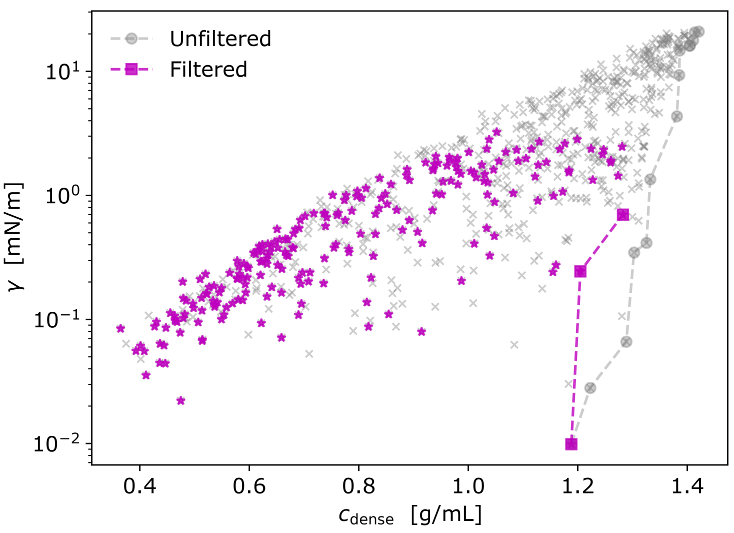


Fig. S2: All data collected from the optimizations shown in Figure 2. The filter applies Waltz[8] and TANGO[9] predictors and imposes a maximum of 5 aromatic residues.

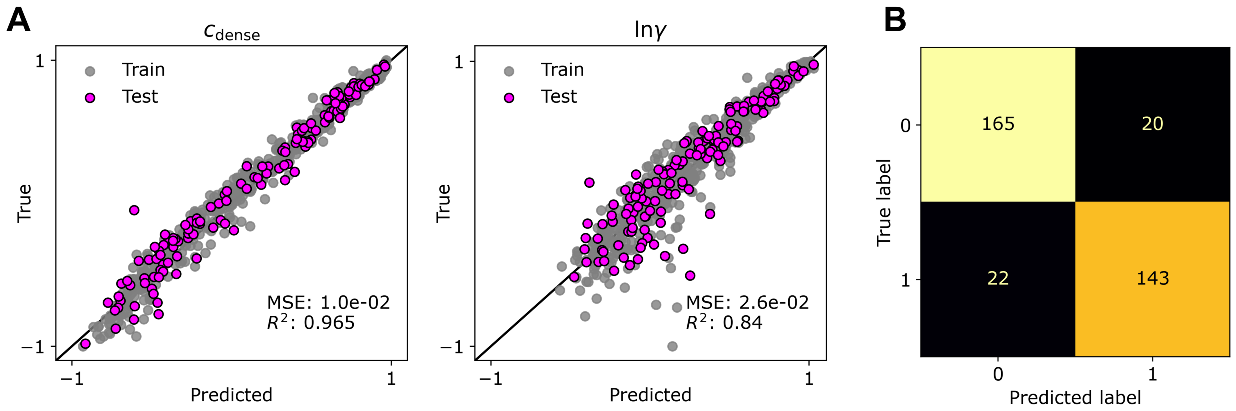


Fig. S3: Performance of the classifier and regressor after the optimizations shown in Figure 2, evaluated on a random 80/20 train-test split.

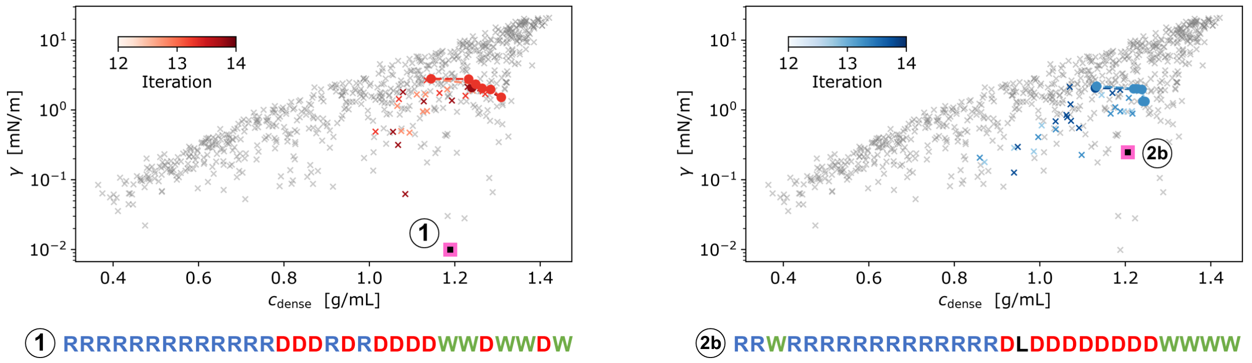


Fig. S4: Optimization progress for constrained optimizations with constant composition in both cases. The left figure is also shown in the main text.

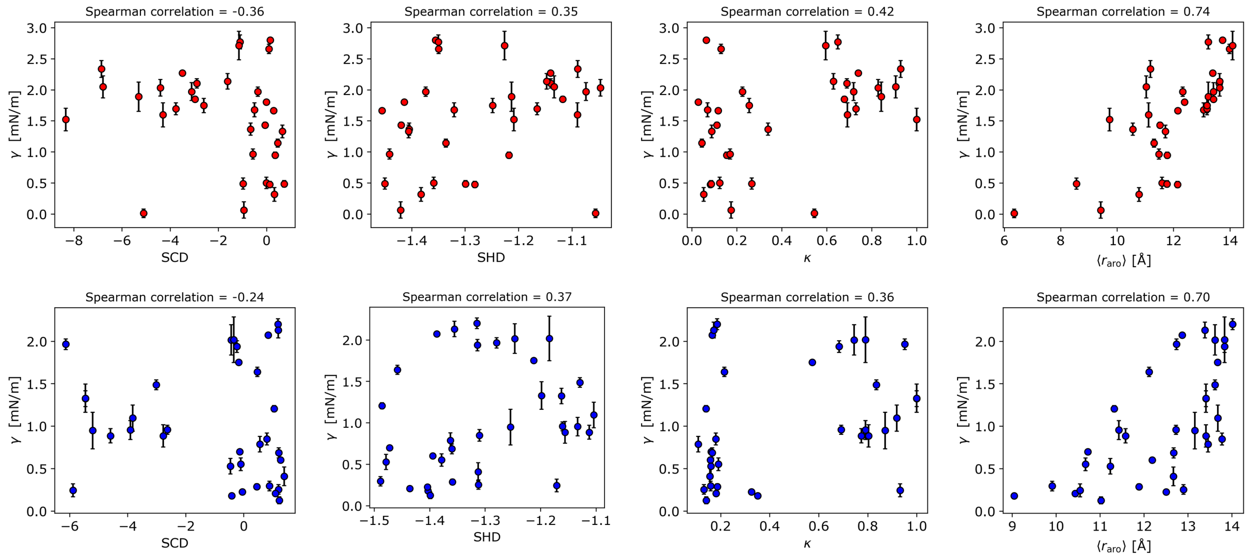


Fig. S5: Analysis of correlations between sequence features and interfacial tension ($\gamma$) at constant sequence composition. Top/Bottom: Optimizations 1/2b, as presented in Figure S4.

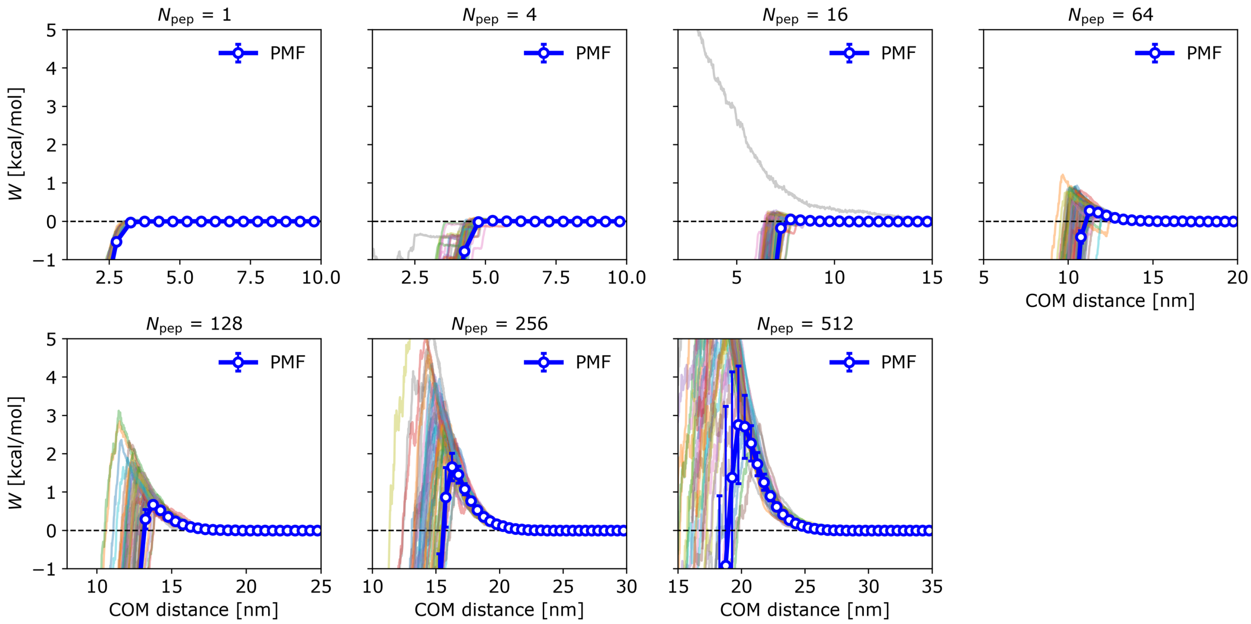


Fig. S6: Quantifying potential barrier for coalescence of different condensate sizes using steered molecular dynamics, as presented in Figure 4. Peptide: low-$\gamma$ peptide 1.

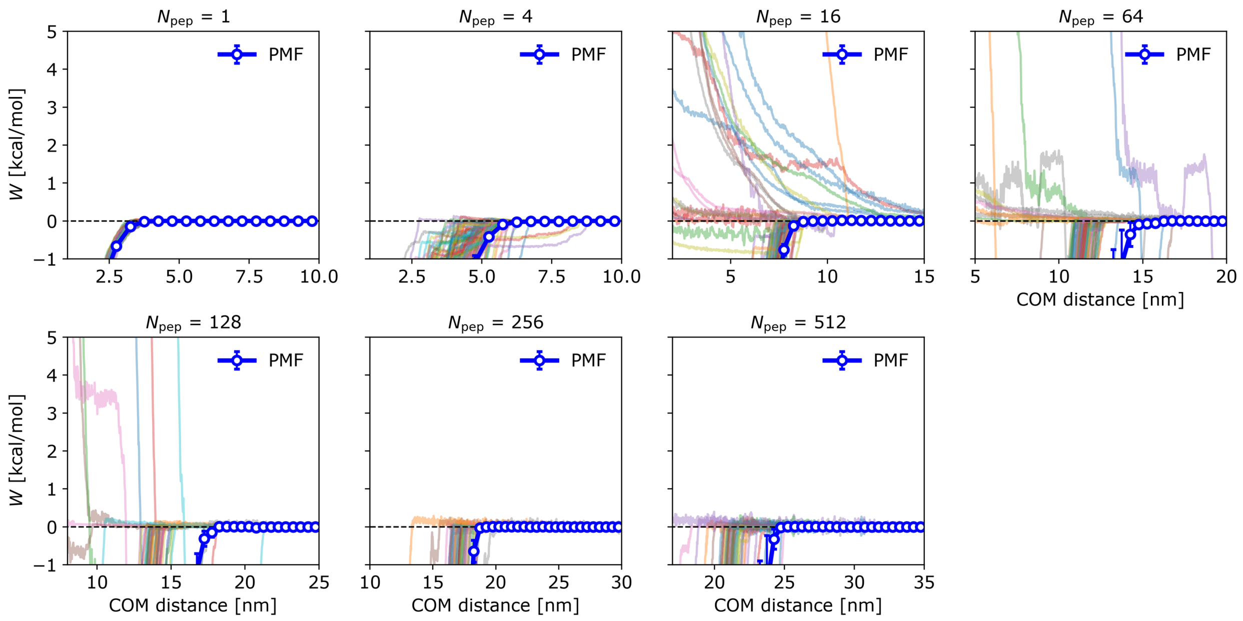


Fig. S7: Quantifying potential barrier for coalescence of different condensate sizes using steered molecular dynamics, as presented in Figure 4. Peptide: high-$\gamma$ peptide 2.

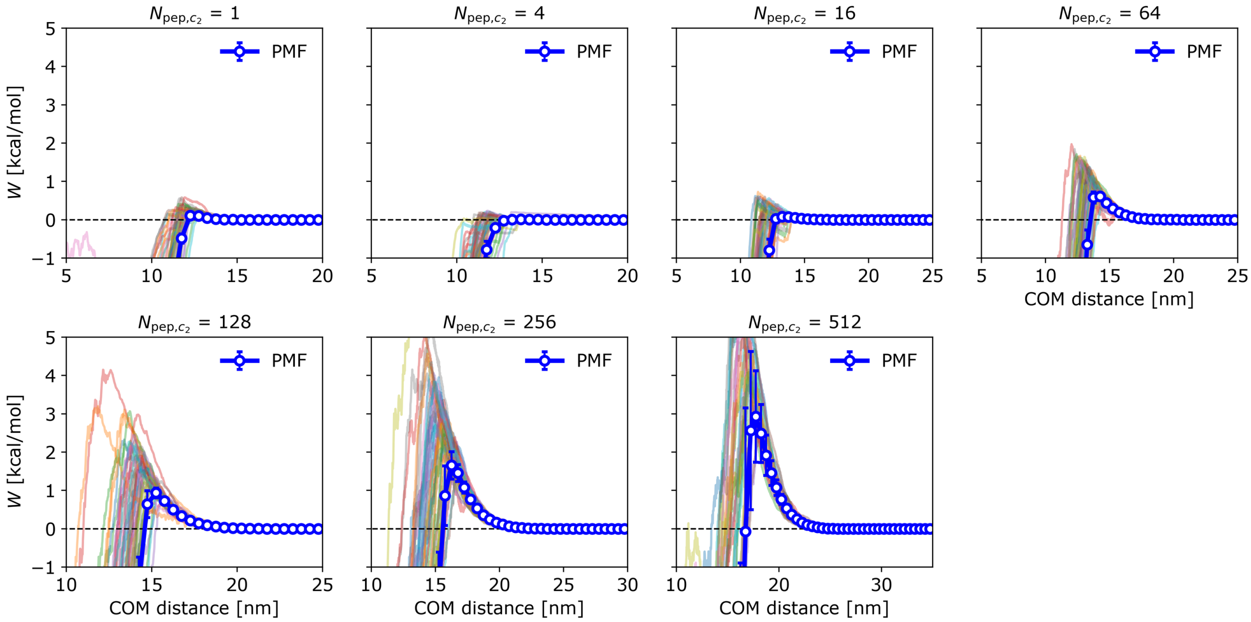


Fig. S8: Quantifying potential barrier for coalescence using steered molecular dynamics with a condensate of 256 peptides and varying the size of the other condensate, as presented in Figure 4. Peptide: low-$\gamma$ peptide 1.

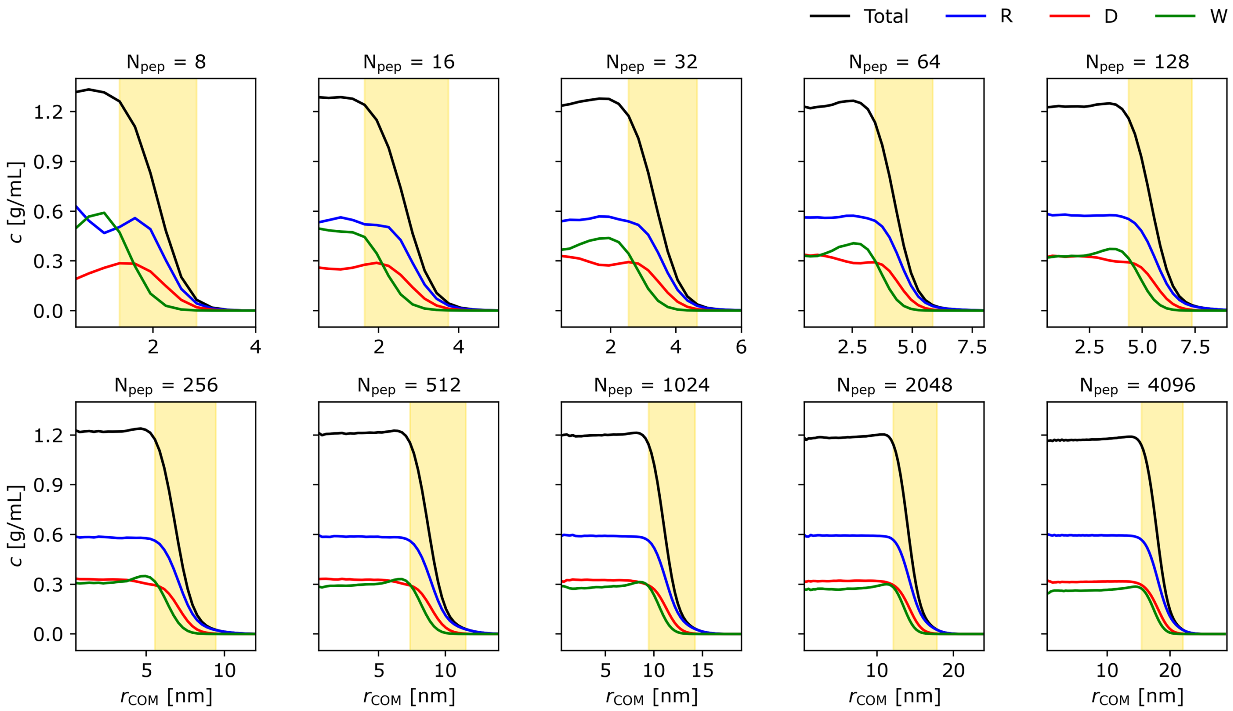


Fig. S9: Concentrations as a function of distance from condensate center-of-mass for varying condensate sizes, as presented in Figure 5. Peptide: low-$\gamma$ peptide 1.

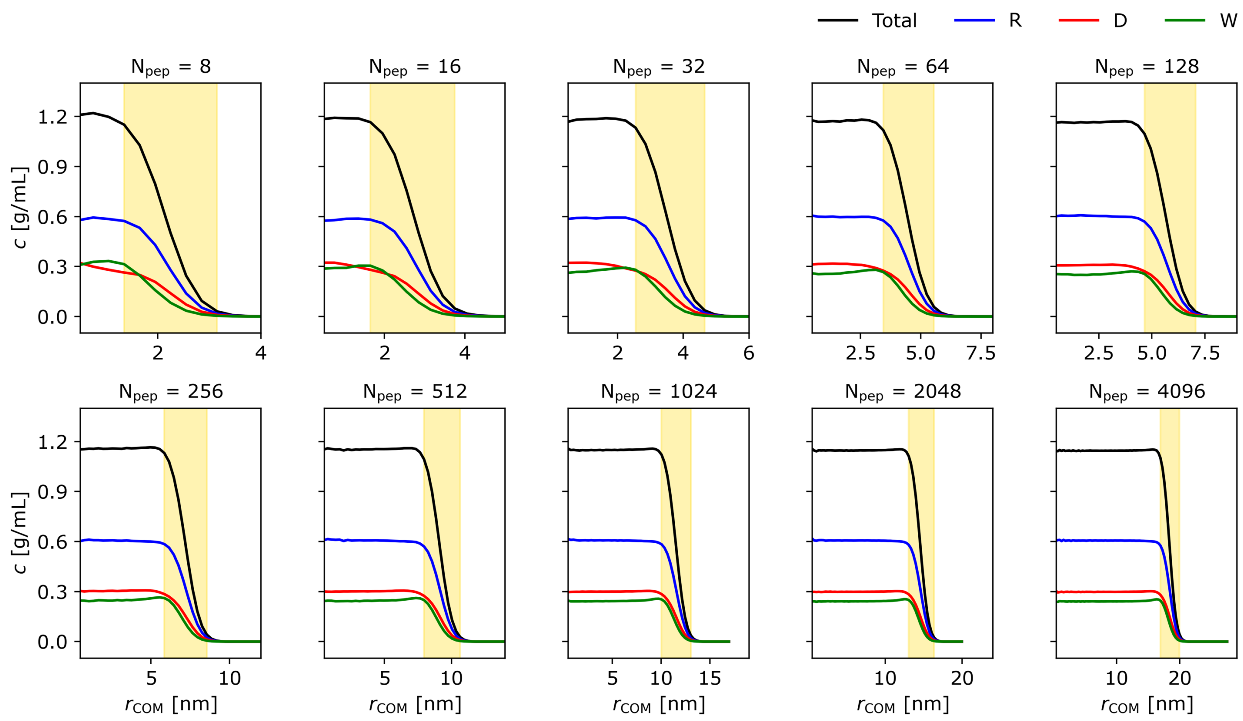


Fig. S10: Concentrations as a function of distance from condensate center-of-mass for varying condensate sizes, as presented in Figure 5. Peptide: high-$\gamma$ peptide 2.

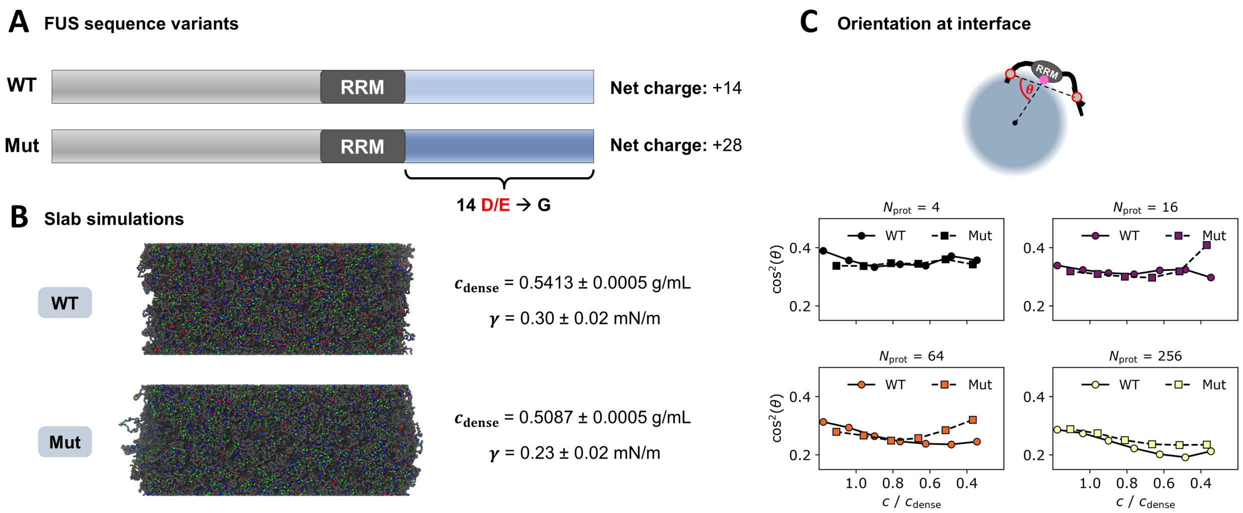


Fig. S11: Analysis of FUS variants[10] using slab coexistence simulations, and molecular orientation analysis with varying condensate sizes analogous to the analysis performed in Figure 5.

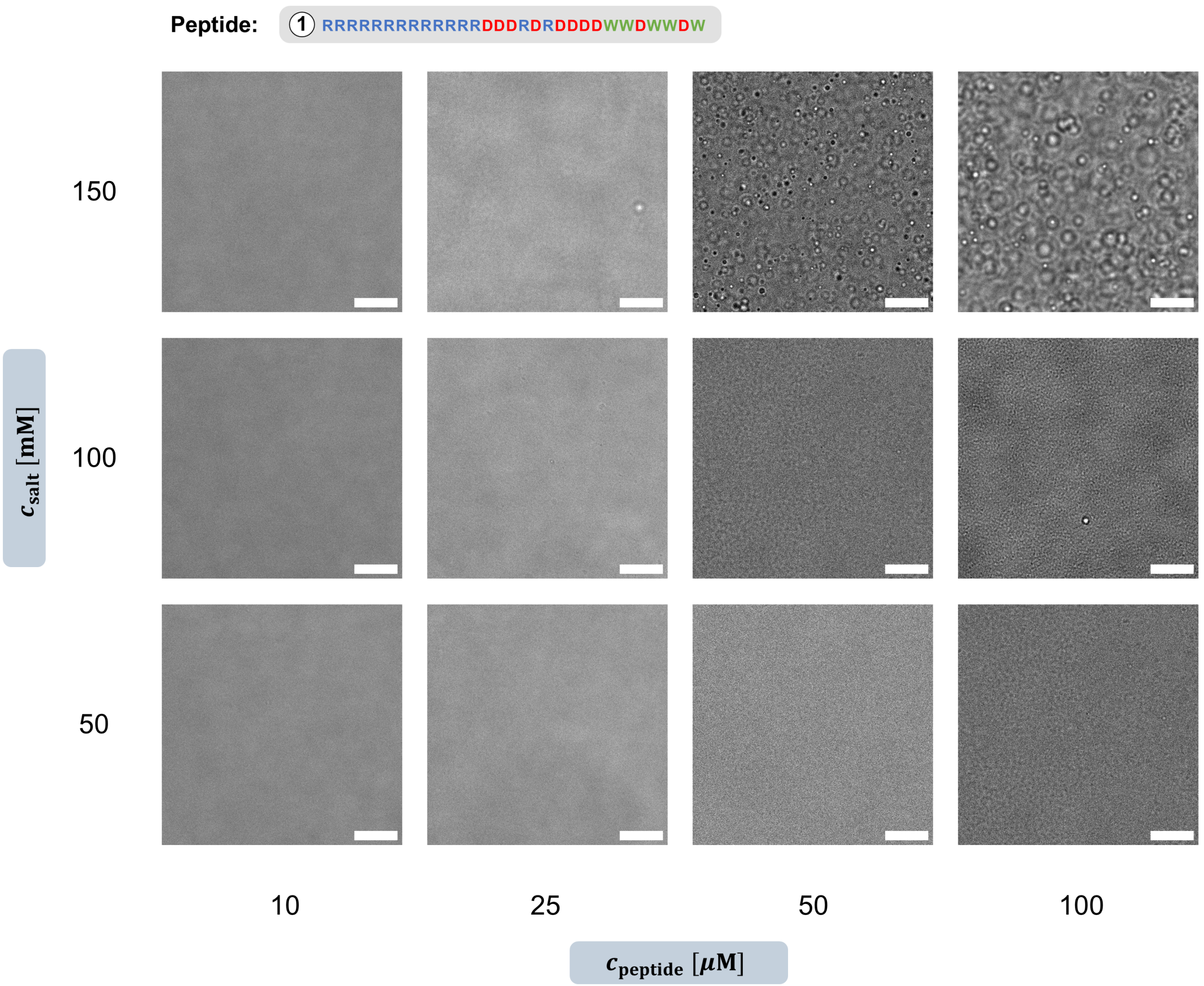


Fig. S12: Bright-field microscopy of peptide 1 solutions at varying concentration and ionic strength. Scale bar: 10 $\mu$m.

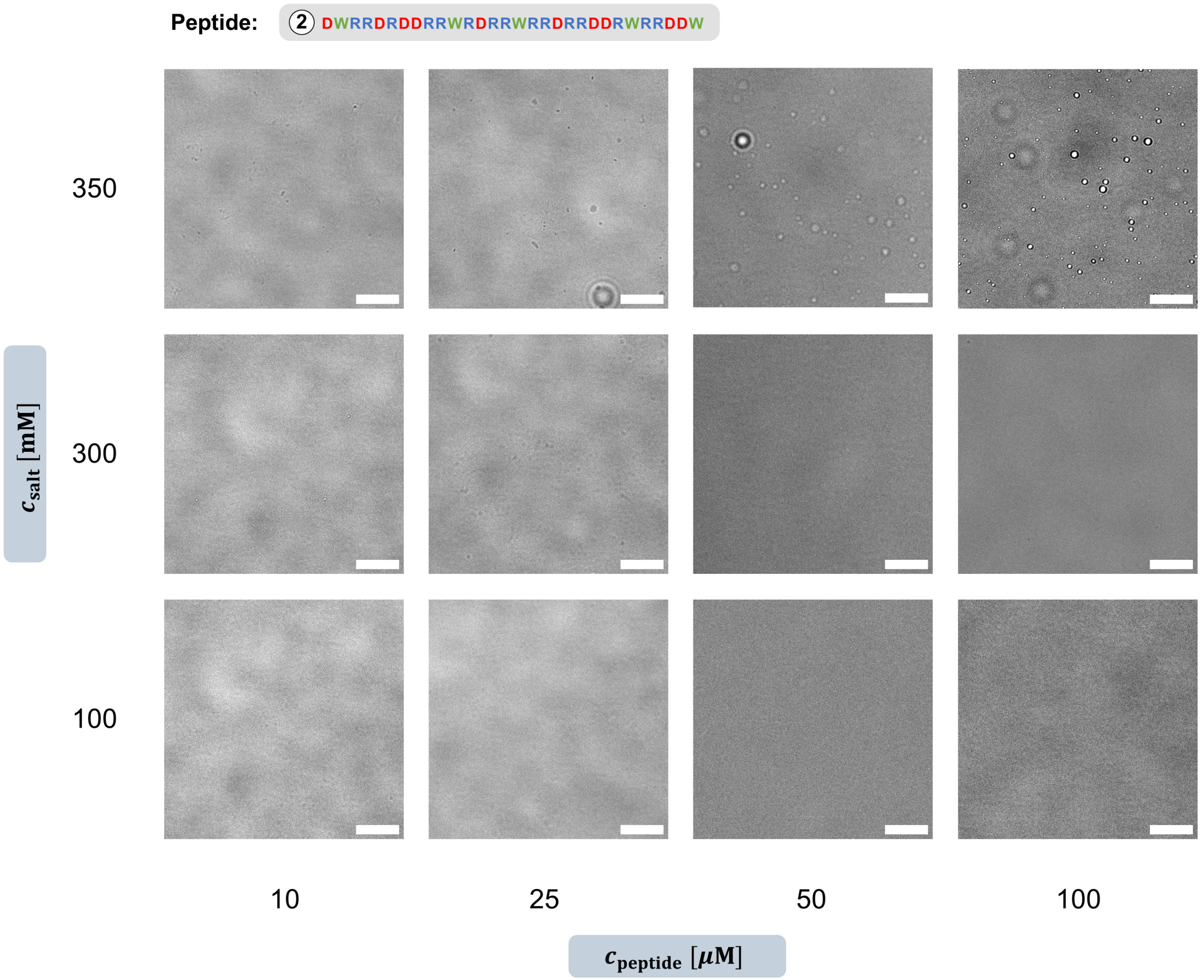


Fig. S13: Bright-field microscopy of peptide 2 solutions at varying concentration and ionic strength. Scale bar: 10 $\mu$m.

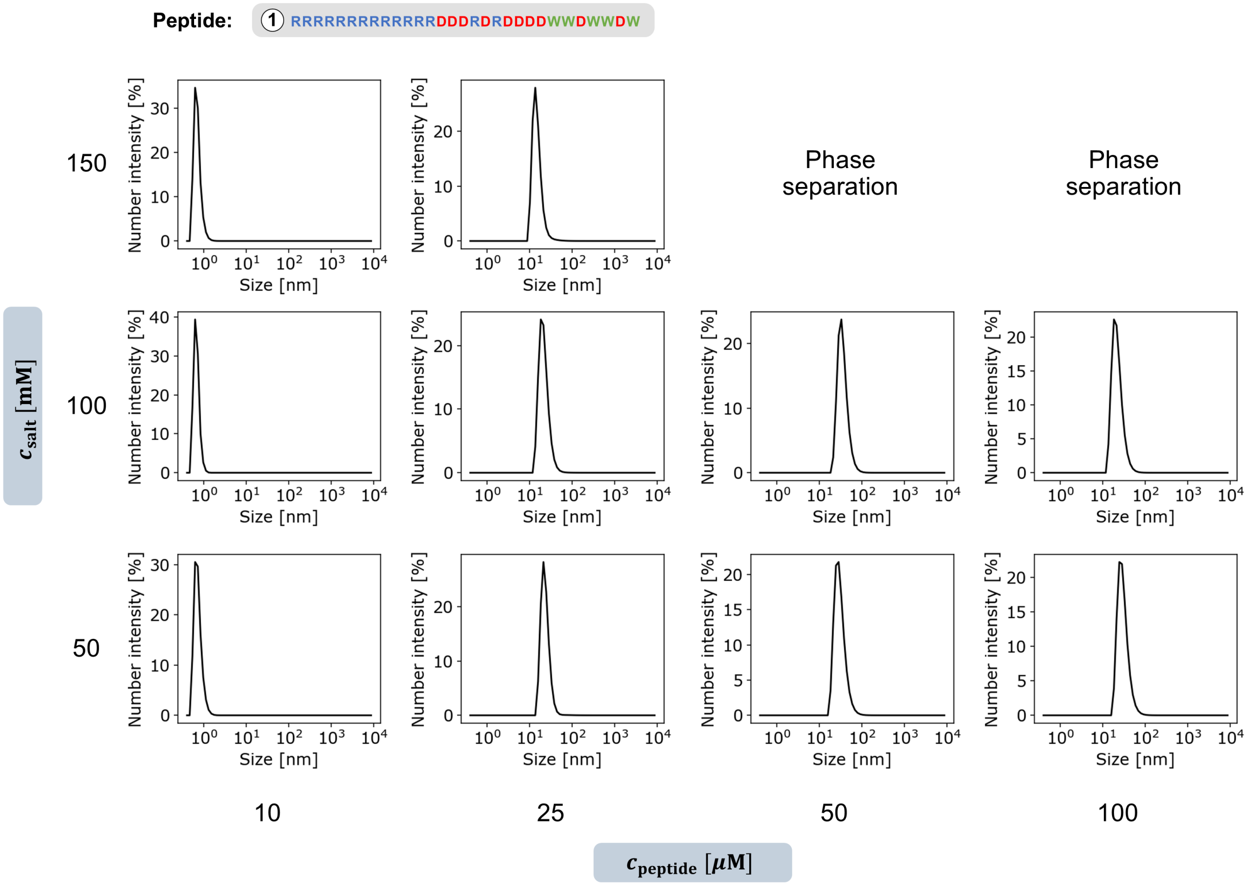


Fig. S14: Dynamic light scattering number intensity profiles for peptide 1 solutions at varying peptide concentration and ionic strength.

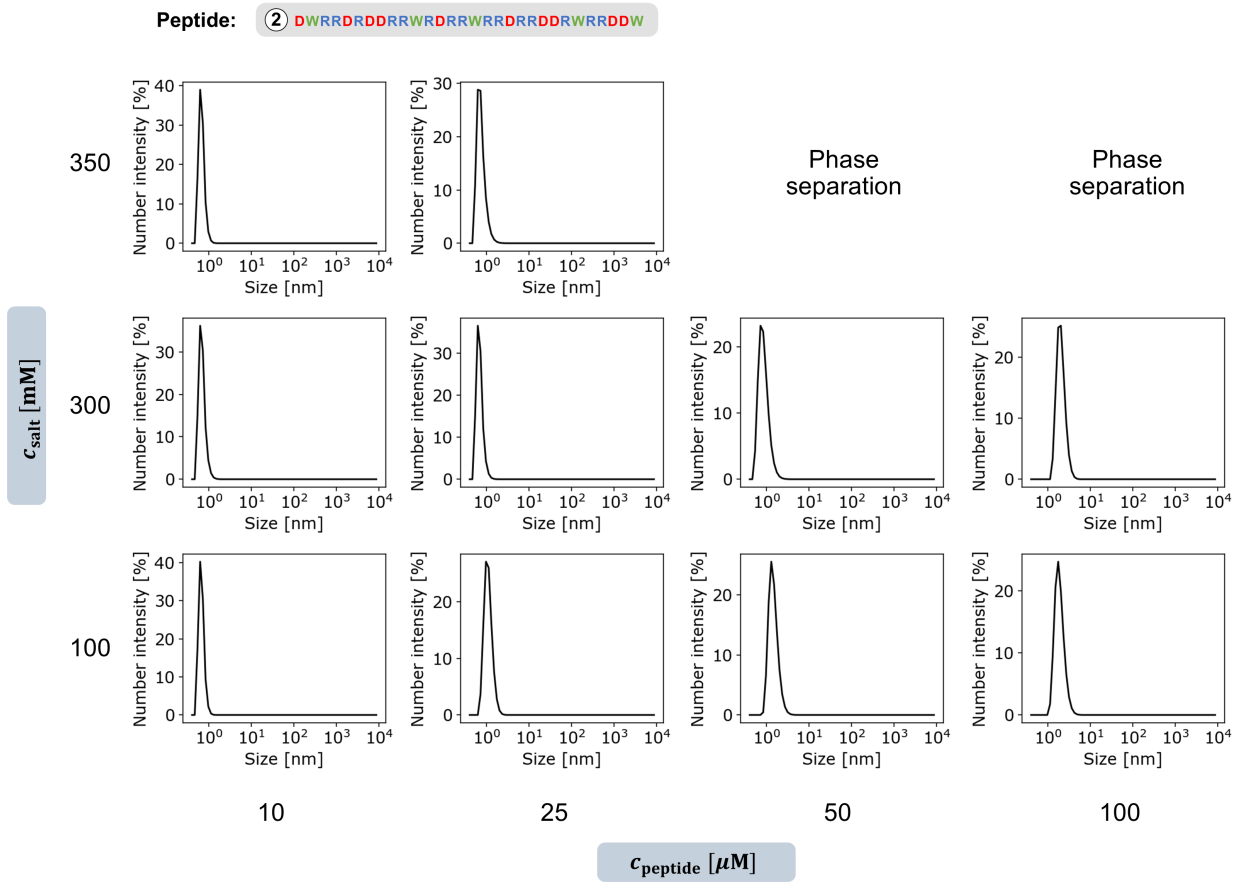


Fig. S15: Dynamic light scattering number intensity profiles for peptide 2 solutions at varying peptide concentration and ionic strength.

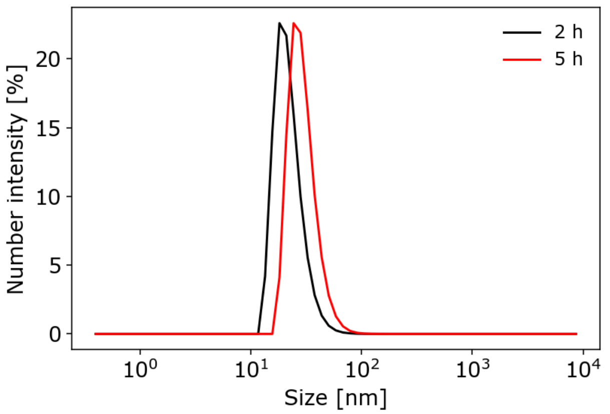


Fig. S16: Nanocondensate time evolution of the dynamic light scattering number intensity of a peptide 1 solution ($c_{peptide} = 100\ \mu M$, $c_{salt} = 100$ mM). A slow shift in the size distribution confirms the metastable character of these species.

Tab. S1: Pareto-optimal sequences after iteration 8. Values are reported as mean ± standard deviation.

| Sequence | $c_{dense}$ [g/mL] | $\gamma$ [mN/m] |
|---|---|---|
| RRRRRWWHWRRMWWRRRWWWWRRRWWWWRR | 1.223 ± 0.059 | 0.03 ± 0.17 |
| RRRRRRRWWRRRWWWWRRRWWWWRRAAWWW | 1.289 ± 0.001 | 0.07 ± 0.47 |
| RRRCRVRRTWRWWWRRSWWWQWRRWWRWWD | 1.303 ± 0.001 | 0.35 ± 0.15 |
| RRWWRRRRRRRWWRWWRRWWRWWRRWWRWW | 1.325 ± 0.001 | 0.41 ± 0.61 |
| RRWRRRRRWRWRWRWWRRWWRWWRRWWRWW | 1.332 ± 0.001 | 1.34 ± 0.86 |
| ERRRRRRRRRWWWRRRWWWWRRRWWWWWDD | 1.381 ± 0.002 | 4.33 ± 0.77 |
| RRRRRRRWRRRWWWWRRRWWWWWEDWWWWD | 1.385 ± 0.000 | 9.34 ± 0.95 |
| RRRWRWRWRRRWWWWRRRWWWWWDDWWWWW | 1.386 ± 0.001 | 14.70 ± 0.32 |
| RRRRRRRRRRRWWWWWDDWWWWWWDDWWWW | 1.405 ± 0.001 | 16.08 ± 4.31 |
| RRRRRRRRRRRWWWWWDDWWWWWWDDWWWW | 1.405 ± 0.001 | 16.08 ± 4.31 |
| RRRRRRRRRRWWWWWDDWWWWWWDDWWWWW | 1.411 ± 0.001 | 17.71 ± 1.16 |
| RRRRRWWRRRWWWWWWDDWWWWWWDDWWWWW | 1.414 ± 0.000 | 20.57 ± 1.39 |
| RRRWWWWRRRWWWWWWDDWWWWWWWDDWWWWW | 1.421 ± 0.002 | 20.87 ± 3.35 |

Tab. S2: Pareto-optimal sequences after iteration 8 and applying Waltz[8] and TANGO[9] filters. Values are reported as mean ± standard deviation.

| Sequence | $c_{dense}$ [g/mL] | $\gamma$ [mN/m] |
|---|---|---|
| RRRRRWWHWRRMWWRRRWWWWRRRWWWWRR | 1.223 ± 0.059 | 0.03 ± 0.17 |
| RRRRRRRWWRRRWWWWRRRWWWWRRAAWWW | 1.289 ± 0.001 | 0.07 ± 0.47 |
| RRWWRRRRRRRWWRWWRRWWRWWRRWWRWW | 1.325 ± 0.001 | 0.41 ± 0.61 |
| RRWRRRRRWRWRWRWWRRWWRWWRRWWRWW | 1.332 ± 0.001 | 1.34 ± 0.86 |
| RRRRWRWWRRWWRWWRRWWRWWRRWWRWRW | 1.349 ± 0.002 | 4.64 ± 0.32 |
| RRRWWRWWRRWWRWWRRWWRWWRRWWRWWR | 1.366 ± 0.002 | 10.34 ± 1.18 |
| WWWWPWWWWRRRWWWWRRRWWWWRRRWWWW | 1.368 ± 0.001 | 18.90 ± 1.49 |

Tab. S3: Pareto-optimal sequences after iteration 11, with maximum 5 aromatic residues. Values are reported as mean ± standard deviation. The last sequence is listed twice, because it was explored twice by the optimization algorithm.

| Sequence | $c_{dense}$ [g/mL] | $\gamma$ [mN/m] |
|---|---|---|
| RRRRRRRRRRRRRDDDRDRDDDDWWDWWDW | 1.188 ± 0.000 | 0.01 ± 0.07 |
| RRWRRRRRRRRRRRRRDLDDDDDDDDWWWW | 1.205 ± 0.000 | 0.24 ± 0.08 |
| RRRRRRRRRRRRRRHEDDDDDDDDDDWWWWW | 1.300 ± 0.000 | 0.67 ± 0.06 |
| RRRRRRRRRRRRRDDDDDDDDDDDDDWWWWW | 1.311 ± 0.000 | 1.25 ± 0.04 |
| RRRRRRRRRRRRRRDDDDDDDDDDDDWWWWW | 1.317 ± 0.000 | 2.03 ± 0.18 |
| RRRRRRRRRRRRRDDDDDDDDDDDDDWWWWW | 1.321 ± 0.001 | 2.48 ± 0.07 |
| RRRRRRRRRRRRRDDDDDDDDDDDDDWWWWW | 1.321 ± 0.001 | 2.50 ± 0.08 |

Tab. S4: Pareto-optimal sequences after iteration 11 and applying Waltz[8] and TANGO[9] filters, with maximum 5 aromatic residues. Values are reported as mean ± standard deviation.

| Sequence | $c_{\text{dense}}$ [g/mL] | $\gamma$ [mN/m] |
|---|---|---|
| RRRRRRRRRRRRRDDDRDRDDDDWWDWWDW | 1.188 ± 0.000 | 0.01 ± 0.07 |
| RRWRRRRRRRRRRRRRRDLDDDDDDDDWWWW | 1.205 ± 0.000 | 0.24 ± 0.08 |
| GRRRRRRRRWWRRWRRRRRWDDDDDDDWEE | 1.283 ± 0.001 | 0.70 ± 0.14 |

Tab. S5: Pareto-optimal sequences after iteration 14, with constant composition of peptide 1 (Figure 2D). In this case, applying Waltz[8] and TANGO[9] filters had no effect. Values are reported as mean ± standard deviation.

| Sequence | $c_{\text{dense}}$ [g/mL] | $\gamma$ [mN/m] |
|---|---|---|
| RRRRRRRRRRRRRRRWWDDDDDDDDDDWWW | 1.309 ± 0.001 | 1.52 ± 0.18 |
| WWDDDDDRRRRRRRRRRRRRRRDDDDWWWD | 1.284 ± 0.001 | 1.97 ± 0.14 |
| RWRRRRRRRRRRRRRRDDDDDDDDDWDWWW | 1.264 ± 0.001 | 2.05 ± 0.18 |
| WRRRRRRRRRRRRRRRDDDDDDDDDWDWWW | 1.249 ± 0.000 | 2.33 ± 0.14 |
| RRWWWRRRRRRDDDDDDDDDDRRRRRRWRW | 1.233 ± 0.000 | 2.77 ± 0.11 |
| DWRRDRDDRRWRDRRWRRDRRDDRWRRDDW | 1.144 ± 0.000 | 2.80 ± 0.04 |

Tab. S6: Pareto-optimal sequences after iteration 14, with constant composition of peptide 2b (Figure 2D). In this case, applying Waltz[8] and TANGO[9] filters had no effect. Values are reported as mean ± standard deviation.

| Sequence | $c_{\text{dense}}$ [g/mL] | $\gamma$ [mN/m] |
|---|---|---|
| LWWRRRRRRRRRRRRRRRDDDDDDDDDWWW | 1.246 ± 0.000 | 1.32 ± 0.09 |
| WWDDDDDDDDDRRRRRRRRRRRRRRRWWWL | 1.241 ± 0.001 | 1.33 ± 0.17 |
| LRWRRRRRRRRRRRRRRWDDDDDDWDDDWW | 1.239 ± 0.000 | 1.96 ± 0.06 |
| WWLWRRRRRRDDDDDDDDDRRRRRRRRWWR | 1.228 ± 0.001 | 2.02 ± 0.18 |
| WWLWRRRRRRRDDDDDDDDDRRRRRRRRWW | 1.219 ± 0.000 | 2.02 ± 0.27 |
| DWRRRRDDWRRRDDWLRRDRRDRRDDWRRW | 1.133 ± 0.001 | 2.20 ± 0.06 |
